# An approach to gene-based testing accounting for dependence of tests among nearby genes

**DOI:** 10.1101/2021.05.24.445494

**Authors:** Ronald Yurko, Kathryn Roeder, Bernie Devlin, Max G’Sell

## Abstract

In genome-wide association studies (GWAS), it has become commonplace to test millions of SNPs for phenotypic association. Gene-based testing can improve power to detect weak signal by reducing multiple testing and pooling signal strength. While such tests account for linkage disequilibrium (LD) structure of SNP alleles within each gene, current approaches do not capture LD of SNPs falling in different nearby genes, which can induce correlation of gene-based test statistics. We introduce an algorithm to account for this correlation. When a gene’s test statistic is independent of others, it is assessed separately; when test statistics for nearby genes are strongly correlated, their SNPs are agglomerated and tested as a locus. To provide insight into SNPs and genes driving association within loci, we develop an interactive visualization tool to explore localized signal. We demonstrate our approach in the context of weakly powered GWAS for autism spectrum disorder, which is contrasted to more highly powered GWAS for schizophrenia and educational attainment. To increase power for these analyses, especially those for autism, we use adaptive *p*-value thresholding (AdaPT), guided by high-dimensional metadata modeled with gradient boosted trees, highlighting when and how it can be most useful. Notably our workflow is based on summary statistics.

## Introduction

More than 3,000 human GWAS have examined over 1,800 diseases and traits, with uneven success in discovering associations [1]. For schizophrenia, for example, 280 discoveries were recently announced, while, for genetically correlated autism spectrum disorder, a handful of loci have been discovered [2]. The difference largely is due to statistical power. To increase power, one might decrease the number of hypotheses tested and thus reduce the threshold for significance. A natural strategy is gene-based testing: SNPs are assigned to genes they occur in or nearby [3]; within this unit, test statistics for SNPs are aggregated; and, finally, significance is judged by the number of genes tested. By focusing tests on genes instead of SNPs dispersed throughout the genome, gene-based testing also has interpretability as an appealing feature. Power can also be enhanced by choosing false discovery rate (FDR) control for significance testing. These two options, gene-based testing and FDR control, are not mutually exclusive. H-MAGMA [4] combines them and also incorporates Hi-C data into its testing scheme. Likewise, when SNPs affect gene expression, these functional SNP-to-gene assignments can be modeled [5].

A related approach is to increase power by incorporating metadata about SNPs or genes in the targeting of multiple testing procedures; selective inference provides approaches to incorporating this information while maintaining valid FDR control. An early approach incorporated metadata directly through the use of *p*-value weights [6]. More recently, in the setting of SNP-based GWAS, we [7] implemented a data-driven approach to determine weights via the adaptive *p*-value thresholding (AdaPT) framework [8]. In brief [7], we improved power for detecting a subset of weakly correlated SNPs by using gradient boosted trees to model potentially-informative metadata, such as known effects of SNPs on gene expression and on genetically correlated traits.

Here we explore the use of AdaPT in the context of gene-based tests for autism (ASD), schizophrenia (SCZ), and educational attainment (EA), placing special emphasis on how it enhances power to detect associations of genes with ASD. To do so, we utilize gene-based testing methods introduced to account for linkage disequilibrium (LD) among SNPs in a gene [9, 10, 11]. LD is not limited by gene boundaries, however. To the contrary, LD among SNPs falling in different genes is commonplace. This compromises the inter-pretability of current gene-based tests, obscuring the meaning of error guarantees with family-wise error rate and FDR controlling procedures. Because of the extensive and heterogeneous LD in the genome, in our prior SNP-based GWAS using AdaPT, we purposely selected quasi-independent SNPs for analysis. By contrast, for gene-based testing, we introduce an agglomerative algorithm to account for LD-induced correlation of test statistics. This algorithm directly groups genes into ‘loci’ for which between-loci test statistic correlation—based on LD—is bounded above. This reduces the set of tests to a collection of weakly correlated genes and loci. Importantly, this agglomerative algorithm can be used in any gene-based testing framework to highlight gene-based tests that are dependent.

We analyze results from three GWAS: ASD[2], SCZ[12] and EA[13]. Using AdaPT guided by loci metadata as our example, we are able to improve power to select ASD-associated genes and LD-defined loci with multiple genes while maintaining finite-sample FDR control. Improvements are more modest for the other phenotypes, due to the high power of their original GWAS. One novel feature of our analyses is that it groups genes into loci when their test statistics are expected to be highly dependent, due to LD. We complement this feature with graphical tools to examine the distribution of association signal within each such locus. The interactive visualization tool we develop uses `R` Shiny [14, 15] and `plotly` [16] for exploring and highlighting biological signals therein.

Our workflow for improving power to select associated genes and LD-defined loci follows (Figure 1): We first introduce the agglomerative algorithm. Then we demonstrate our approach for detecting associations with in the AdaPT framework. For metadata, we use eQTL data from cortical tissue samples, gene co-expression networks [17], and GWAS results for the other phenotypes, which is motivated by the observation that all three are genetically correlated[18]. Relationships of metadata to gene-based statistic are uncovered by use of gradient boosted trees. We separate our results into two categories, technical features related to gene-based testing and implementation of AdaPT in this setting, and association results for these phenotypes and their implications, with special emphasis on ASD.

**Figure 1:**
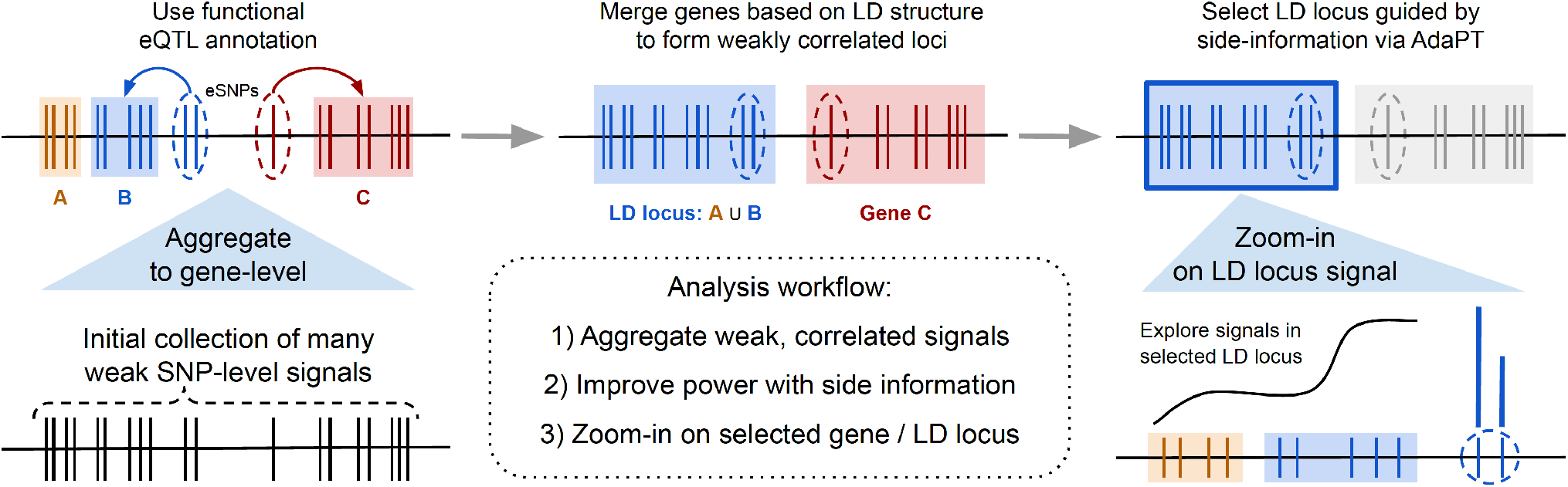
Schematic of our workflow to improve power with metadata and highlight interesting signals while adjusting for LD structure.

## Methods

### SNP-to-gene assignment and correlation between gene-level tests

We consider two different approaches for assigning *n* SNPs to a set of genes *G*. Following common practice, we use genomic location only for “Positional” SNP assignment: SNP *i* is assigned to gene *g* if its genomic position is within gene *g*’s start and end positions 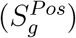. In some published analyses, the start and end positions are expanded slightly to include presumed regulatory regions, although we do not do so here. As an alternative approach, recognizing that some SNPs have documented effects on genes, such as eQTL effects, we use Positional SNP assignment and include in the set cis-eQTL SNP-gene pairs, which we dub eSNPs. We denote the collection of eSNPs for gene *g* as 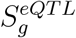 and call this “Positional + eSNPs” SNP assignment.

After assigning SNPs to genes, each gene’s vector of SNP-level *z* statistics ***z**_g_* is modeled as multivariate normal (Gaussian) with mean 0 and LD-induced covariance **Σ**_*g*_. Following common practice [9], we assume **Σ**_*g*_ is known using correlations from reference genotype data, specifically 503 individuals from the 1000 Genomes EUR sample as the reference data [19]. Gene-level testing frameworks [3, 9, 10] combine SNP-level signals into gene-level test statistics *T_g_* while accounting for **Σ**_*g*_, the LD among SNPs in a gene, but ignore the LD between SNPs involved in gene-level test statistics for nearby genes *g* and *g*′.

Consider the quadratic gene-level test statistics, *T_g_* = ***z_g_**^T^ **z_g_***, featured in *VEGAS* [9, 10] and MAGMA (v1.08). Let *S_g_* and *S_g′_* be the two sets of SNPs for *g* and *g*′, respectively. Under the null model, the induced covariance between the quadratic test statistics for these two sets of SNPs can be computed as,

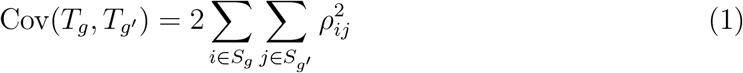

where *ρ_ij_* is the correlation between SNPs *i* and *j* based on reference data. Using the fact that the variance for a single set of SNPs *S_g_* is,

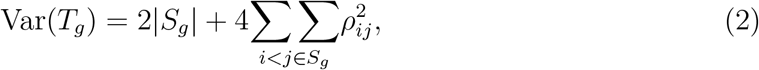

we can compute the induced correlation between the quadratic test statistics for two sets of SNPs 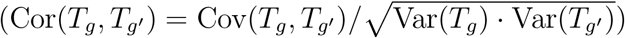.

For nearby gene sets *g* and *g*′, these correlations can be quite strong. This can confound the interpretability—and even meaning—of the guarantees from multiple testing procedures. To avoid these issues, we will modify the construction of our gene sets to bound the pairwise correlation between the resulting gene sets.

### Agglomerative LD loci testing

To account for substantially correlated test statistics, we introduce an agglomerative procedure to group highly correlated genes that are within *w* megabases (Mb) into sets of genes we refer to as LD loci or simply loci. Given an LD threshold *r*^2^, we apply the following procedure to a set of genes *G* within a chromosome:

1. Compute Cor(*T_g_, T_g′_*) for all pairs of genes, *g, g*′ ∈ *G*, within *w* Mb of each other (using Equations 1 and 2).
2. Repeat the following until (Cor(*T_g_, T_g′_*))^2^ *< r*^2^ for all remaining pairs of genes/loci in *G*:

- Find genes/loci 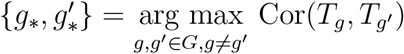,
- Merge 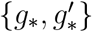 into locus *g_LD_*,
- Update 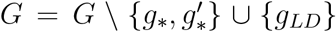, and compute Cor(*T_gLD_, T_g′_*) for all *g*′ ∈ *G* within *w* Mb of *g_LD_*.

This is essentially agglomerative hierarchical clustering, but with a linkage determined by the LD-based correlation structure of the test statistics. We compute the quadratic test statistic, *T_g_*, for each remaining gene/locus *g* ∈ *G*.

Because the resulting distribution *T_g_* does not have a known closed-form solution, we use a Monte Carlo based approach for computing the *p*-value *p_g_* for the gene/locus to test the null hypothesis that its *n_g_*-dimensional vector of SNP-level *z* statistics are not associated with trait status. We generate *B* draws of null, *n_g_*-dimensional Gaussian random variables, 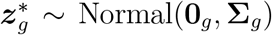. A quadratic test statistic 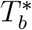 is calculated for each of the *b* ∈ [*B*] draws, resulting in an empirical *p*-value:

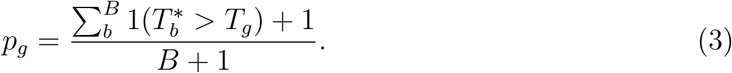

### Overview of GWAS data and eQTL sources

Our investigation focuses on reported GWAS *z* statistics, {*z_i_, i* = 1*, …, n*} measuring SNP-level association with ASD [2], SCZ [12] and EA [13]. For GWAS results of one phenotype, we explore SNP-level association statistics from the other two GWAS as potential sources of metadata due to previous evidence of their genetic correlation [18]. We consider *n* = 5, 238, 256 SNPs whose alleles could be aligned across all three phenotypes and with minor allele frequency (MAF) *>* 0.05 based on the 1000 Genomes EUR sample reference data [19]. Also, for these SNPs, their hg19 variant locations could be converted to GRCh38 using the LiftOver utility from the UCSC Genome Browser (http://genome.ucsc.edu/). Probably due to a smaller sample size, ASD has lower power: 18,381 cases and 27,969 controls, in comparison to SCZ with 33,426 cases and 32,541 controls, and EA with ≈ 1.1 million subjects (Supplementary Figure 1). Because we focus on detecting associations for ASD, a neurodevelopmental disorder, we leverage two different sources of cortical tissue to identify eSNPs for functional SNP-to-gene assignment. The first source of eSNPs was obtained from the BrainVar study of dorsolateral prefrontal cortex from 176 individuals sampled across a developmental span [20]. We identified 151,491 cis-eQTL SNP-gene pairs meeting BH *α* ≤ 0.05 for at least one of the three sample sets: prenatal (112 individuals), postnatal (60 individuals), as well as across the complete study. This corresponds to 123,664 eSNPs associated with 6,660 genes, with 85% of the eSNPs associated with one unique cis-eQTL gene pairing.

The second source is adult cortical tissue cis-eQTLs from the Genotype-Tissue Expression (GTEx) V7 project dataset [21]. Instead of using eQTLs as reported by GTEx, to be consistent with the BrainVar eQTL definition, we identified 414,405 cis-eQTL SNP-gene pairs meeting BH *α* ≤ 0.05 for either *Frontal Cortex BA9* or *Anterior cingulate cortex BA24* samples based on the tissue specific files for all SNP-gene associations available at gtexportal.org. This resulted in 313,316 GTEx eSNPs associated with 9,012 genes, where 78% of the eSNPs are associated with one gene. However we observe an overlap of 55,313 cis-eQTL SNP-gene pairs with BrainVar, culminating in 510,583 unique cis-eQTL SNP-gene pairs with 370,749 eSNPs associated with 12,854 genes across the union of BrainVar and GTEx sources.

### GENCODE version

We use GENCODE v21 [22] for our list of genes with their respective start and end positions based on genome assembly version GRCh38. This matches the version used in the BrainVar study, but differs from GTEx, which is based on v19. When identifying GTEx eQTLs, we removed 187 genes from GENCODE v19 that do not match Ensembl IDs in v21. This provides us with an initial list of *G* = 57, 005 candidate genes to potentially assign SNPs to, based on either positional or functional eSNP status.

### Metadata

For each gene/locus *g*, we create a vector of metadata *x_g_* collected independently of *p_g_*. This process is completed in the same manner for both the *Positional* and *Positional + eSNPs* collections. First, we consider the number of SNPs assigned to a gene/locus, *n_g_* = |*S_g_*|, which can be viewed as statistical information relevant to the power of the quadratic test statistic. Additionally, we include one-sided *z* statistics, i.e., *z_g_* = Φ^−1^(1 − *p_g_*), constructed using the gene/locus-level *p*-values from independent GWAS results. For our target phenotype ASD we use SCZ and EA GWAS *z* statistics, 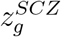 and 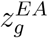, while for SCZ (EA) we use 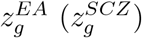 and 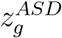 as metadata.

Given the set of eSNPs 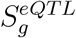 associated with single genes or genes in LD locus *g*, we summarize the expression level as the average absolute eQTL slope in a relevant source to obtain 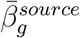 for five sources: three BrainVar developmental periods (pre-, post-, and complete) and two adult GTEx cortical regions. Furthermore, we account for weighted gene co-expression network analysis (WGCNA) [17] modules by creating two sets of indicators, one set for the twenty modules reported in the BrainVar study and another for eight modules constructed using the GTEx cortical tissue samples. For simplicity, we also construct indicators denoting if gene/loci is not included in any of the modules.

We also include additional context about the gene/loci. Indicator variables determine four GENCODE biotypes: protein coding, antisense, long non-coding RNA, and other. Using gnomAD v2.1.1 [23], we associate with each gene its loss-of-function observed / expected upper fraction (LOEUF) value, which indicates the gene’s tolerance to loss-of-function. Because a lower LOEUF scores indicate strong selection against loss-of-function, we include the minimum LOEUF across all genes in an LD locus in our vector of metadata *x_g_*.

### AdaPT implementation

Given a collection of gene/locus-level *p*-values and metadata, (*p_g_, x_g_*)_*g*∈*G*_, we apply AdaPT to select a subset of discoveries with FDR control at target level *α* = 0.05. AdaPT is guaranteed finite-sample FDR control under the assumption of independent null *p*-values, and was demonstrated to maintain control in weak, positive correlated scenarios [7], such as the one considered here.

We incorporate metadata from the feature space 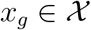 using gradient boosted trees, implemented in XGBoost [24], to separately model the non-null probability and effect size (see *Supplementary information* for details). This gives us flexibility to include many potentially useful covariates without being overly concerned about the functional form with which they enter the model or their marginal utility. However, overfitting in this context will lead to a loss of power. To find appropriate settings for the gradient boosted trees (number of trees, learning rate, and maximum depth), we first “tune” AdaPT’s performance with synthetic SCZ *p*-values that are aligned with the ASD *p*-values in the following manner:

1. Sort SCZ and ASD *p*-values: 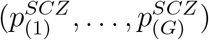 and 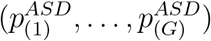
2. Replace SCZ with ASD *p*-values by matching order, i.e., replace 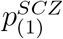 with 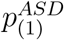, 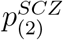 with 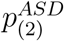, …

This transforms the SCZ signal to match the weaker signal in ASD. We then proceed to apply AdaPT using these synthetic SCZ *p*-values to find candidate settings which yield the highest number of synthetic SCZ discoveries at FDR level *α* = 0.05. Finally, for each phenotype and positional assignment, we use two cross-validation steps within AdaPT [7] to select from among these candidate settings to generate our AdaPT: XGBoost results using the `adapt_xgboost_cv()` function in the `adaptMT R` package (available at https://github.com/ryurko/adaptMT).

### Kernel smoothing localization

Following the selection of interesting genes/loci, researchers may be interested in “zooming” in on localized signals at the gene and/or SNP-level. For a selected gene/locus *g\** and its corresponding set of SNPs *S_g*_*, we smooth over the squared *z* statistics of the locus’ positional SNPs 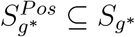, given their genomic positions (BP) using the Nadaraya–Watson estimator:

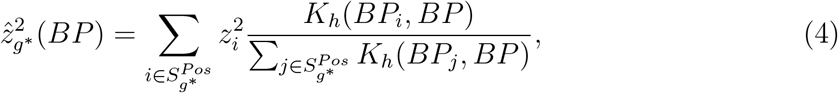

in which *K_h_* is a one-dimensional Gaussian kernel with bandwidth *h* selected separately for each gene/locus using generalized cross-validation (as implemented in the `np` package [25]). We then provide the option to display any subset of eSNPs 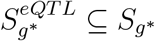 separately as bars with their heights indicating individual SNP-level signal. This separation is due to the presence of intergenic eSNPs, however any eSNPs that are also positionally assigned to genes, 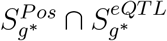, are included in the positional smoothing in Equation 4.

## Results

### Assigning SNPs to genes and generating LD loci

We assign SNPs to genes using the two approaches: *Positional*, which assigns 2,779,780 SNPs to 40,581 genes; and *Positional + eSNPs*, which includes an additional 109,042 intergenic cortical tissue eSNPs resulting in 2,888,822 SNPs assigned to 41,301 genes. Next, we generate genes/loci based on the LD-induced correlation of gene-based test statistics using the agglomerative algorithm with window size *w* = 6 Mb and with one of three thresholds for *r*^2^, 0.25, 0.50, 0.75. The number of independent genes/loci decreases substantially as the threshold becomes more strict for both SNP assignment types (Figure 2). Even a relatively high threshold of *r*^2^ = 0.75 reduces the number of *Positional + eSNPs* (*Positional*) gene/locus tests from 41,301 (40,581) to 37,522 (37,114). We report the conservative threshold *r*^2^ = 0.25 in the body of the manuscript (see *Supplementary information* for results with *r*^2^ ∈ {0.50, 0.75}). Due to the conservative threshold we combine 17,915 genes to form 4,136 LD loci for *Positional + eSNPs* and 16,625 genes to form 3,985 LD loci for the *Positional* approaches. Over 75% of these loci contain five or fewer genes while the largest is a chromosome 11 locus that groups over sixty genes, most of which encode olfactory receptors. Combined with the 23,386 and 23,956 individual genes that were not merged, this results in 27,522 and 27,941 genes/loci for testing. The reduction of independent testing units highlights the correlation among genes that is often ignored in gene-based testing. We then compute the gene/locus quadratic test statistics and *p*-values for each phenotype using the Monte Carlo-based approach in Equation 3 with *B* equal to two million simulations.

**Figure 2:**
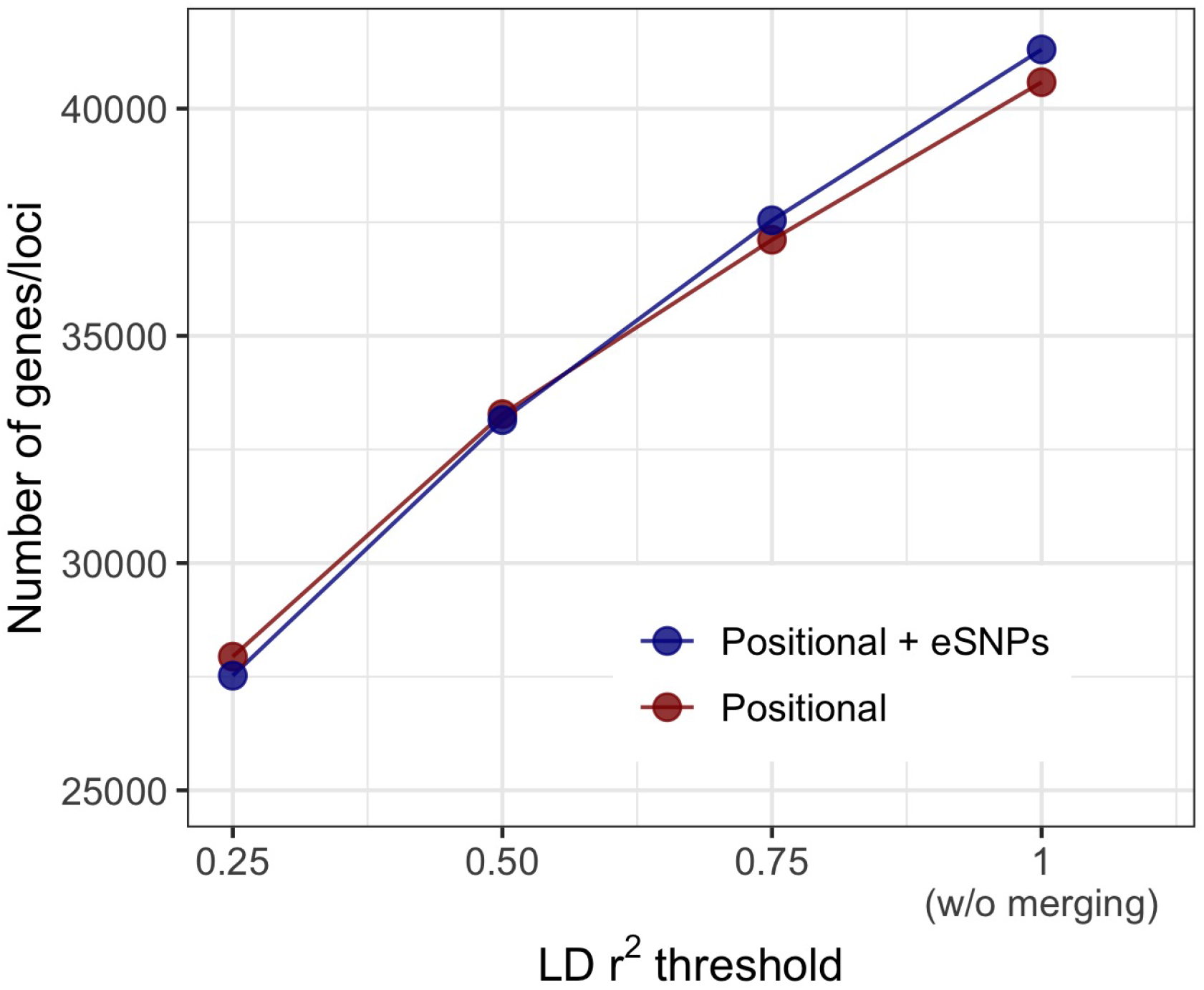
Comparison of the number of genes/loci following our agglomerative algorithm over a range of values for the induced LD threshold *r*^2^ ∈ {0.25, 0.50, 0.75} by positional type (in color). The initial number of genes is provided for reference (corresponding to *r*^2^ = 1).

### AdaPT models and results

To generate AdaPT: XGBoost results, we first tune the procedure based on synthetic SCZ *p*-values, which mimic the distribution of ASD p-values, to find optimal XGBoost settings. To avoid over-fitting, we consider shallow trees with maximum depth ∈ {1, 2}, while searching over the number of trees *P* ∈ 100*, …, * 450 by increments of fifty and the learning rate *η* ∈ 0.03*, …, * 0.06 by increments of 0.01. To analyze the real *p*-values for each phenotype and thereby select associated genes/loci, then, these top setting combinations (Supplementary Table 1) were used in the AdaPT cross-validation steps and while targeting FDR control at *α* = 0.05.

To find genes/loci associated with each phenotype, our AdaPT: XGBoost implementation fits a mixture model that requires two parameters – the probability of association and the effect of each gene/locus on the phenotype – and these are estimated separately for each step of the algorithm. Variables in the metadata inform on each of these parameters to different degrees; see *Supplementary information* and Supplementary Table 2 for details on measuring variable importance. For the *Positional + eSNPs* results, the number of SNPs per gene/locus and z-statistics for at least one genetically correlated trait are important predictors for all three phenotypes (Figure 3). SCZ and EA, in contrast to ASD, display increased importance for LOEUF and membership in a WGCNA module constructed from the GTEx cortical tissue samples (Figure 3) Lower LOEUF values, which indicate lower tolerance to loss-of-function, were more likely to be associated with SCZ and EA. We observe similar patterns of variable importance for the *Positional* results (Supplementary Figure 2).

**Figure 3:**
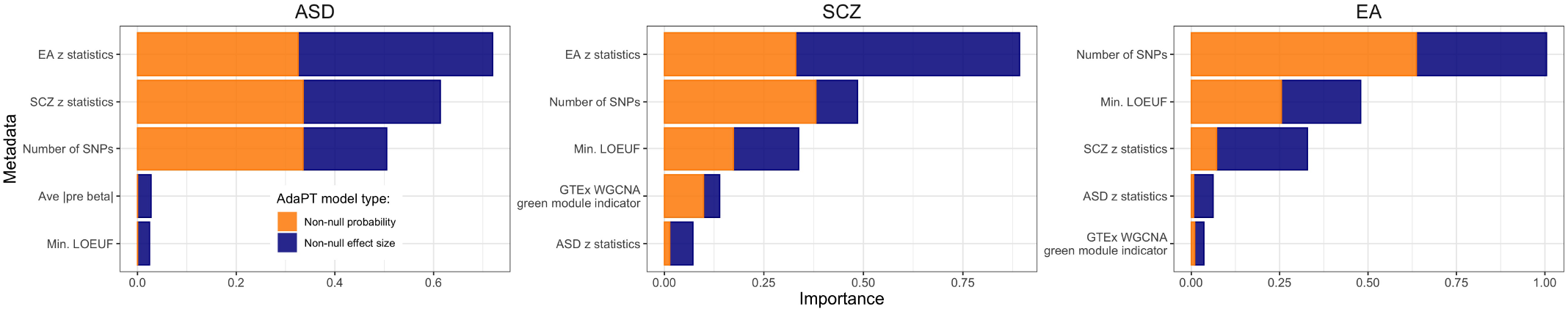
The most important variables (top five) predicting association with phenotype as ranked by XGBoost for the *Positional + eSNPs* assignment of SNPs. Variables are sorted in order of importance, while the color of the bars denote the separate parameters for the AdaPT implementation: probability of association (orange) and of non-zero effect size (blue).

Comparing the number of genes/loci selected by AdaPT to baseline results of intercept-only versions of AdaPT and BH, there is a clear gain in gene/locus discovery by accounting for metadata through the AdaPT: XGBoost implementation, regardless of phenotype and SNP-to-gene assignment approach (Figure 4A, Supplementary Table 3, see also *Supplementary information*, Supplementary Figures 3 and 4 for results with LD threshold of *r*^2^ ∈ {0.50, 0.75}). Unsurprisingly, the number of associated genes/loci is much larger for SCZ and EA than for ASD, likely due to the lower power of the original ASD GWAS. For *Positional* ASD, we see that the intercept-only version of AdaPT fails to select any genes/loci due to the weak signal without inclusion of metadata (Figure 4A).

**Figure 4:**
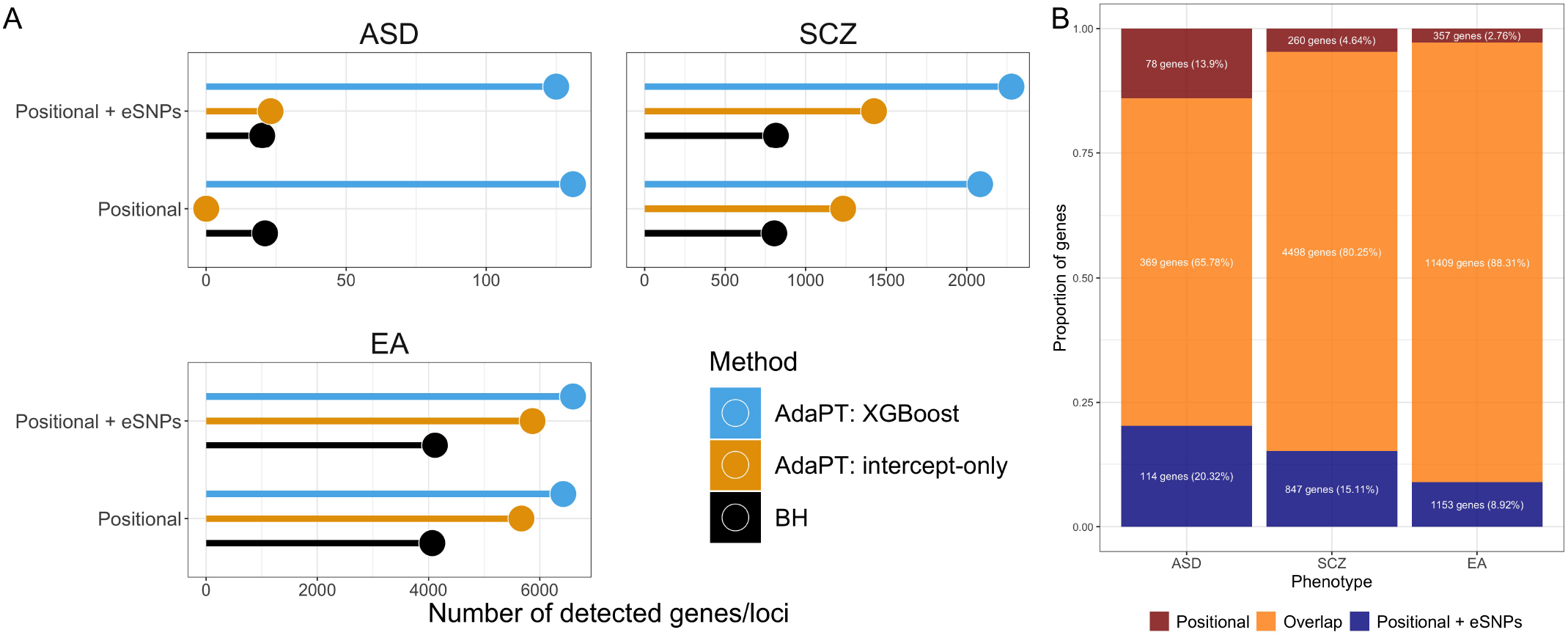
*(A)* Comparison of the number of selected genes/loci at FDR level *α* = 0.05 for each phenotype by positional assignment. AdaPT: XGBoost results are presented in comparison to BH and AdaPT: intercept-only baselines which do not account for metadata. *(B)* Comparison of the proportion of implicated genes that overlap between the two types of positional assignment results, based on the AdaPT: XGBoost results for each phenotype.

### Comparison of phenotypic results

For ASD, analysis of *Positional + eSNPs* identifies 483 genes, of which 405 cluster in 47 loci and 78 are unclustered, whereas analysis of *Positional* SNPs alone yields 447 genes, of which 370 cluster in 54 loci (Supplementary Table 3). A substantial portion of these genes overlap (Figure 4B). While similar patterns emerge for SCZ and EA, the ratio of unclustered to clustered genes increases with increasing number of genes/loci detected: 0.193 for ASD, 0.414 for SCZ, and 0.681 for EA. This presumably reflects greater power to detect small effects of a SNP on phenotype with larger sample size: decay of this signal tends to cause it to fall below the threshold of detection for SNPs in nearby genes. In contrast, the proportion of genes uniquely identified when eSNP information is included is substantially higher for ASD than it is for SCZ or EA (Figure 4B), again likely due to lower power for the ASD sample.

As expected, for all three phenotypes, the number of unclustered genes increases with increasing threshold *r*^2^ (Supplementary Table 4). If, however, we assume that signal should be sparsely distributed across the genome, then the sum of LD loci and unclustered genes, for *r*^2^ = 0.25, should be a reasonable estimate of genes associated given the current data. This translates into 125 genes for ASD, 2,277 for SCZ, and 6,598 for EA, and correspondingly 0.30, 5.51, and 15.98% of the total 41,301 genes analyzed (including protein-coding and non-coding genes). For SCZ and EA, the total number of associated genes per chromosome declines strongly and linearly with chromosome size (Supplementary Figures 5–7), which itself is correlated with the total number of genes per chromosome. This pattern, while more variable, is also seen in a breakdown of genes into protein-coding and other non-coding types (Supplementary Figure 6). For ASD, however, an unusually large number of genes are associated on chromosomes 3, 8, 15, and 17, relative to chromosome size, and the associated gene counts show only a modest relationship with chromosome size (Supplementary Figures 5–7), presumably due to lower power and resulting lower number of associated genes per chromosome.

### Exploring signal in selected genes/loci

LD loci, as we define them, are expected to exhibit correlated association signal. Nonetheless, the signal is unlikely to be distributed evenly across the locus, instead in many instances it will be concentrated near one or more SNPs generating the signal, depending on the LD pattern in the locus. The same is true for signal in non-clustered genes. To interactively explore localized signal within genes/loci, we developed an LD locus zoom application using `R` Shiny [14, 15] and `plotly` [16] (available here: https://ron-yurko.shinyapps.io/ld_locus_zoom/). This tool displays the gene/locus of interest, represents genes by their location therein, and highlights the association signal by kernel smoothing for positional SNP signals, including interpolation.

#### 3.7 Mb deletion region in chromosome 17

One of the associated loci, roughly 500 Kb, falls in the well-known 3.7 Mb 17p11.2 deletion/duplication region associated with Smith-Magenis syndrome (deletion, OMIM:182290) [26] and Potocki-Lupski syndrome (duplication, OMIM:610883) [27]. The associated LD locus displays overlapping genes and signals from eSNPS and positional SNPs for ASD, SCZ, and EA (Figure 5A). For reference, the gray dotted line denotes a point-wise 95^*th*^ percentile of 1,000 simulations for squared null Gaussian random variables, 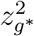 where *z_g*_* ~ Normal(**0**_*g\**_, **Σ**_*g\**_), given the LD structure **Σ**_*g\**_ of the selected LD locus *g\**. Notably, a single gene in the locus has been associated with all three phenotypes to various degrees, *RAI1* [28]. Known to be dosage-sensitive, both deletion and duplication of a single copy of the gene is sufficient to elevate the likelihood of ASD substantially and to diminish cognitive function strongly [28]. Curiously, only the smoothed signal for EA association peaks over *RAI1*, whereas it peaks over *TOM1L1* for ASD and further proximal for SCZ. The peak association for EA over *RAI1* also coincides with the SNP having the lowest p-value for association with this phenotype: this index SNP rs11655029 has p-value 2.84 × 10^−9^ and falls in an intron of *RAI1*. While *RAI1* is a prime candidate for the target gene in this locus for ASD, due to prior evidence, the peak signal located over *TOM1L2* is intriguing, especially because its index SNP rs4244599, while not strongly associated (p-value = 0.0018), is an eQTL for *TOM1L2*. The index SNP for SCZ, rs8082590, has p-value 6.02 × 10^−8^, close to the traditional threshold for GWAS, and coincides roughly with its smoothed signal for association over *GID4*.

**Figure 5:**
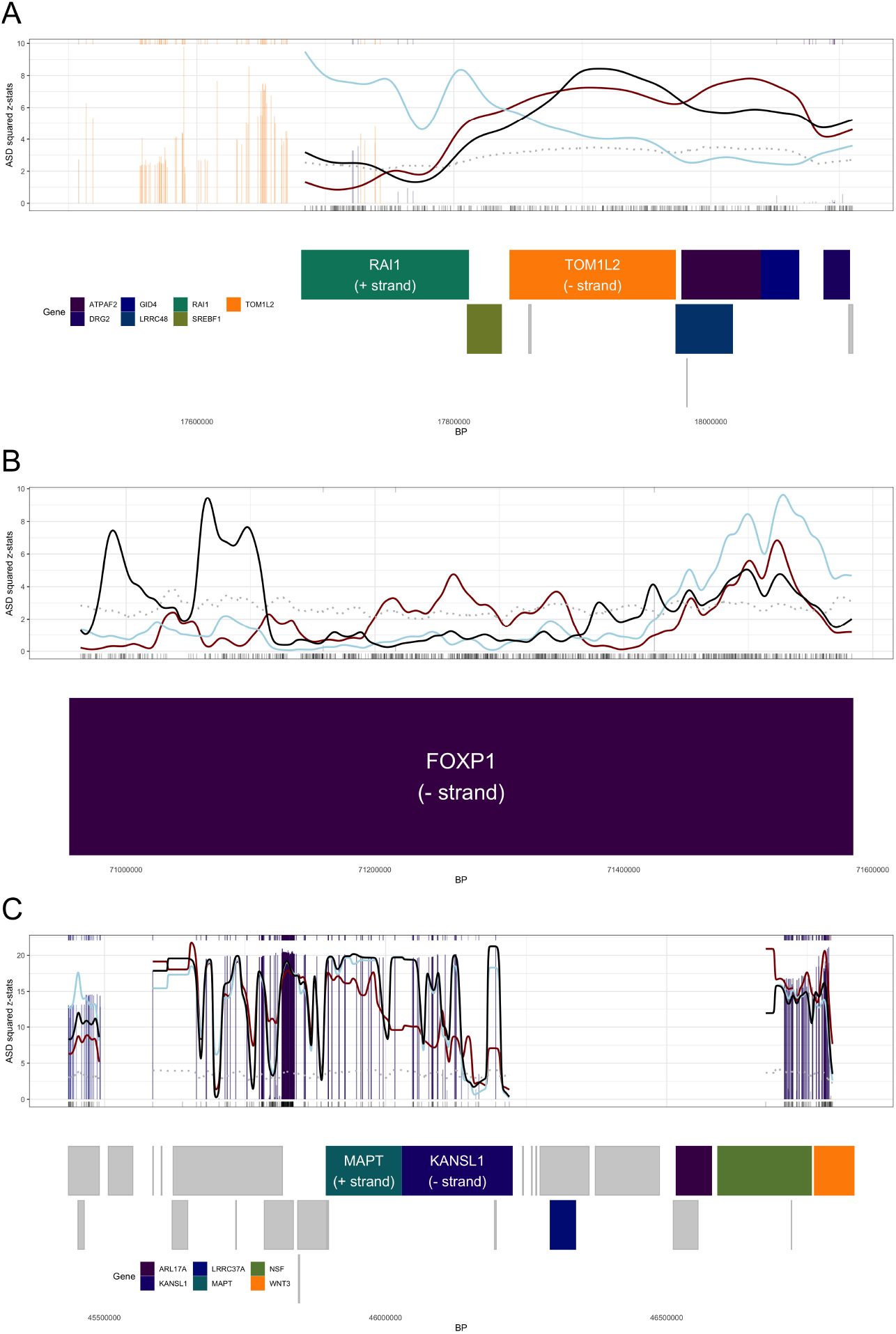
Gene/locus zoom displays for *(A)* locus falling in the chromosome 17 3.7 Mb Smith-Magenis syndrome region, *(B) FOXP1* in chromosome 3, and *(C)* locus in chromosome 17 1 Mb inversion region. *(A)*-*(C)* The genes located within the locus are represented in rectangles denoting their respective start and end positions below the smooth display, arranged by position and size to prevent overlapping. The gene/locus level kernel smoothing of ASD signal (black line) is displayed along with SCZ (red line) and EA (blue line) kernel smoothing signal (both are normalized to appear on the same signal scale as ASD). The gray dotted line denotes a point-wise 95^*th*^ percentile of 1,000 simulations for null simulations, and vertical bars correspond to eSNP signals (colored by their associated genes). A subset of genes are highlighted with colors in *(A)* and *(B)*, with other genes represented by gray rectangles. Additionally, individual genes are labeled in *(A)*: *RAI1* and *TOM1L2*, *(B)*: *FOXP1*, and *(C)*: *MAPT* and *KANSL1*, with gene direction (+ indicates left to right, - indicates right to left).

#### *FOXP1* in chromosome 3

An unclustered protein-coding gene on chromosome 3 highlights a set of associated SNPs in *FOXP1*, which has previously been strongly associated with intellectual disability, language impairment, and ASD [29] (Figure 5B). The smoothed association signal for ASD falls near the gene’s 3’ end whereas, for SCZ and EA, it falls close to the gene’s 5’ region. It is possible that variation associated with ASD is regulating *FOXP1*’s expression quite differently than that for SCZ and EA. The index SNP for EA is GWAS-significant and the one for SCZ approaches it (rs55736314 and rs4677597 with p-values 1.63 × 10^−16^ and 3.87 × 10^−7^ respectively) and both fall closer to the 5’ region of *FOXP1*. For ASD, its index SNP rs7616330 carries a more modest signal (p-value 2.26 × 10^−4^). Compared to the complete gene, it falls toward the 3’ end, but it also falls quite close to the 5’ start site of certain transcripts, such as ENST00000650387.1. Given the prominent role that *FOXP1* has in ASD and cognitive function, we speculate that the differential location of signal for SCZ/EA versus ASD could trace to different transcripts and regulation of their expression.

#### 1 Mb inversion region in chromosome 17

Another locus of interest is a 1.5 Mb region of chromosome 17, namely 17q21 (Figure 5C). This region of the genome is well known in human genetics because it comprises a 1.5 Mb inversion polymorphism [30, 31] and the inversion alleles, actually haplotypes, have been associated with a wide variety of neurodegenerative disorders, including Progressive Supranuclear Palsy [32], corticobasal degeneration [33], frontotemporal dementia [34], and other tauopathies [35]. In this locus, altered *MAPT* is well known to affect risk for late-life neurodegenerative disorders. Moreover, the inversion itself inhibits recombination, rendering alleles at SNPs across this region in high LD. Indeed, the complexity of this region inspires the interactive features of our application (conveyed in Supplementary Figures 8 and 9) with subsets of genes and their associated eSNPs. Our results suggest variation in the region is associated with all three phenotypes. The index SNP for EA exceeds the standard threshold for single SNP significance (rs74998289, p-value 1.31×10^−17^), while for ASD it approaches it (rs12942300, p-value 2.06×10^−6^). Variation in the region has been previously implicated in ASD susceptibility [36]; more recently, *KANSL1* expression has been implicated in cognitive function and ASD [37]. While the smoothed association signals for ASD and EA show a peak over this gene, signal is distributed across many genes in this locus and gene-based analysis is unlikely to pinpoint any gene therein. A clue to the driver or drivers of association comes by comparison of panels A-C in Figure 5. In the inversion region (Figure 5C), eSNP signals are noticeably more enriched compared to the other two loci and indeed the index SNP for EA is an eSNP for *KANSL1*. As highlighted by the display (Figure 5C), however, careful statistical and molecular analyses will be required to pinpoint what variation and what genes influence each of the three phenotypes at this locus.

### Enrichment analysis

A primary motivation for gene-based analyses is to garner insight into the biological mechanisms underlying the phenotype by evaluating the set of genes associated with it. A standard approach infers these mechanisms by gene-set enrichment analysis, which we will implement in two ways. First, we will use the FUMA GENE2FUNC tool [38] for gene ontology (GO) enrichment analysis of the AdaPT: XGBoost genes/loci. This analysis has the advantage of searching through myriad functional sets of genes in an agnostic fashion. Still, this could also be viewed as a disadvantage if, *a priori*, certain biological functions are likely to affect the phenotype. For example, for all three phenotypes analyzed here, rare variation has already provided evidence linking synaptic, epigenetic, and transcription factor genes to them. Moreover, for ASD there exists a substantial set of genes implicated in risk by studies of rare variation. For these reasons, we implement a second gene-set enrichment analysis, specifically GSEA [39], which is a tool for testing if different sets of genes are enriched in a ranked gene list. We perform GSEA at the gene/locus-level ranked by their one-sided *z* statistics, using five different sets of genes/loci to test for enrichment at the top of the phenotype-specific ranked list: (1) brain expressed, (2) synaptic, (3) epigenetic, (4) transcription factors, and (5) 102 ASD risk genes identified based on *de novo* and case-control variation [40] (see *Supplementary information* and Supplementary Table 5 for details on compilation of gene lists).

Another reason to use both two approaches is that they allow us to handle a confounder, gene size, in different ways. In our analyses, gene/locus size (and the number of SNPs therein) is a predictor of association: larger genes/loci are more likely to be associated (Supplementary Figure 10). So enrichment analysis should control for gene size in some way. One approach is to contrast the associated set of genes with control genes after matching on gene size. We will use this approach in the FUMA analyses (see *Supplementary information* for details). However, a substantial portion of brain-expressed genes are synaptic, which are known to be among the largest genes in the genome. We reasoned that if these phenotypes were influenced by variation altering function of synaptic genes—which rare variant studies suggest they are [41, 40, 42, 43]—then matching on gene size will tend to match associated synaptic genes with other synaptic genes, thereby over-matching and lowering the power to detect synaptic association. For this reason, in the GSEA analysis, we first remove the effect of gene size on the association *z* statistics by regressing them on gene size, then entering the residual *z*′ values into the GSEA analysis.

Rather than include all genes in the associated loci for the FUMA enrichment analysis, we used the kernel smoothing results to identify *signal* genes (see *Supplementary information*). For example, this reduces the number of ASD *Positional + eSNPs* genes from 483 to 464 *signal* genes. After matching for gene size, no GO terms show enrichment for associated ASD genes. Genes associated with SCZ display GO enrichment: 698 terms in biological processes, 163 terms in cellular components, and 108 in molecular function. The EA genes display enrichment for 114 terms in biological processes, 64 in cellular components, and 27 in molecular function (Supplementary Table 3). Using REVIGO [44] to summarize these terms, they highlight neuron projection, synaptic function, cell adhesion, cell cycle, chromosome organization, and many more for both SCZ and EA (Supplementary Figures 11 and 12).

GSEA analysis of five gene sets (brain expressed, synaptic, epigenetic, transcription factor, and ASD risk) finds ASD implicated genes enriched for two gene sets, synaptic and epigenetic genes (Figure 6, Supplementary Figure 13), whereas for SCZ and EA, all five gene sets are enriched (Supplementary Table 3).

**Figure 6:**
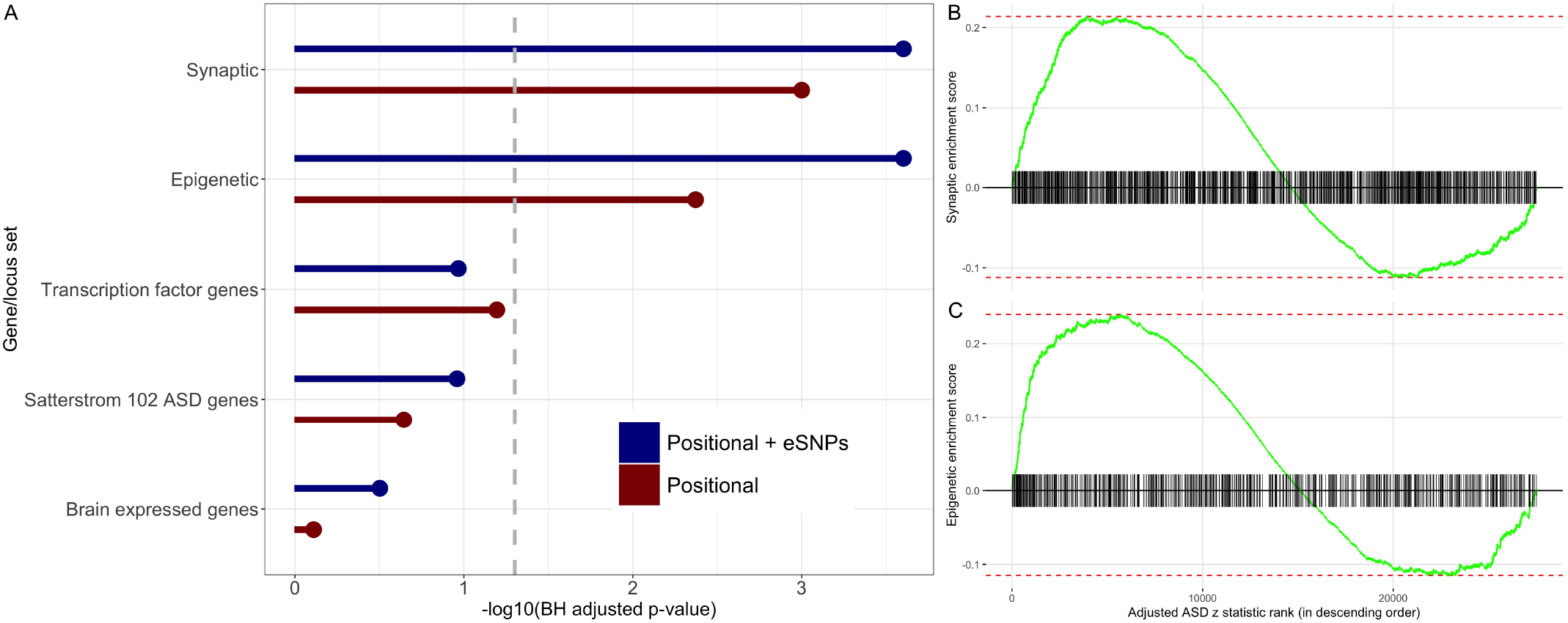
Enrichment analysis of genes/loci ranked by one-sided ASD z statistics, adjusted for size, by positional assignment. *(A)* BH-adjusted GSEA *p*-values are displayed on the -log_1_0 scale, with FDR target level *α* = 0.05 indicated by dashed gray line. Enrichment score for *(B)* synaptic and *(C)* epigenetic genes/loci with tick marks denoting gene/locus rank based on *Positional + eSNPs* ASD z statistics, adjusted for size, in descending order.

## Conclusion

GWAS studies have identified myriad associations between SNPs and human phenotypes. With few exceptions, these associations are weak. One of the goals in this study is to improve power to detect weak associations through gene-based tests, while also accounting for LD among SNPs in different genes, which can induce correlations among tests for different genes. Although such correlation can elevate the false positive rate and obscure the interpretation of gene-based test results, most published studies have ignored it. To account for LD, we introduce a workflow (Figure 1) to aggregate genes into loci if the expected dependence of their test statistics exceeds a pre-specified threshold. This approach produces a notable reduction in the number of genes tested: nearly 4,000 are aggregated even for a tolerant threshold for correlation of test statistics (aggregate if *r*^2^ *>* 0.75; Figure 2). With a collection of uncorrelated or only weakly correlated genes/loci, any gene-based test can be applied. To improve power, we adopt the FDR control framework for hypothesis testing and a particular implementation of it, adaptive p-value thresholding (AdaPT). AdaPT has the potential to increase power over the classical Benjamini–Hochberg (BH) procedure for FDR control by accounting for covariates, which collectively we call metadata, to inform on which genes are likely to be true positives even though their test statistics failed to cross the BH threshold for significance. Importantly, like BH, AdaPT also maintains finite-sample FDR control.

We applied AdaPT to data from three phenotypes, ASD, SCZ, and EA, all of which are genetically correlated. Notably, the ASD GWAS was weakly-powered compared to the well-powered GWAS for SCZ and EA. Although AdaPT increased power for every phenotype and setting (Figure 4A, Supplementary Figures 3 and 4), relative to BH, the largest improvement was achieved for ASD. Gene-based tests that incorporated the impact of SNPs on gene expression also increased power of gene discovery (Figure 4B, Supplementary Figure 5). Reflecting the genetic correlation, summary statistics for other phenotypes were always useful predictors for the AdaPT model (Figure 3, Supplementary Figure 2). Gene conservation also played an important role, as did the size of the gene/locus (Figure 3, Supplementary Figure 10).

That gene/locus size was a useful predictor for associated genes is of limited biological and genetic interest; however, it generates an interpretative challenge when the set of associated genes are evaluated for functional relevance, such as by GO enrichment analysis. One approach to enrichment analysis would match associated genes with control genes by size and we did this in our GO analyses. Such an approach, however, could be conservative. For example, a substantial portion of brain-expressed genes are synaptic, they tend to be large, and synaptic genes likely play a role in all three phenotypes analyzed here [41, 40, 42, 43]. Yet, matching on gene size will tend to contrast associated synaptic genes with other synaptic genes, lowering power to detect their enrichment. For this reason, we also took a different approach to enrichment analysis. Specifically, we regressed the association statistics on size, then entered the residual value into a GSEA analysis. The end result, for ASD, is that synaptic genes were not enriched when genes were matched before GO analysis, while they were enriched in the GSEA analysis (Figure 6, Supplementary Figure 13). The same is true for a set of epigenetic genes, which include chromatin readers, remodelers, and so on. Both results agree well with previous rare variant studies [41, 40]. On the other hand, for the well powered SCZ and EA studies, such enrichment emerges regardless of the way gene size is controlled (Supplementary Figures 11 and 12).

Here we demonstrated the gain in power of evaluating gene-based association statistics using AdaPT, guided by metadata, to detect genes affecting three specific phenotypes, ASD, SCZ, and EA. As our results show, the greatest gain in power is achieved when the underlying study has intermediate power, neither too high nor too low. Power gains will be modest for highly powered studies and absent for very weakly powered studies. We believe that AdaPT, guided by metadata, can be applied to a wide variety of omics problems, although it will undoubtedly require some adaptation for the specific problem to be solved. The advantage of doing so is twofold, increased power and increased interpretability. We are especially interested in the latter, as a means of expediting our understanding of the etiology of human diseases, disorders, and other phenotypes.

Although gene-based association draws attention to an important functional unit (or units for locus-based association), it does not inform on what variation drives the association. We developed an interactive visualization tool (https://ron-yurko.shinyapps.io/ld_locus_zoom/) for exploring the localization of association signal within associated genes/loci and generating hypotheses about mechanisms of action (Figure 5, Supplementary Figures 8 and 9). For example, using this tool to explore the association of *FOXP1* [29] reveals that the pattern of association in the gene is different for ASD versus EA and SCZ (Figure 5B) and suggests that different patterns of gene expression could be important for these phenotypes. Another example is the 500Kb associated locus within the 17p11.2 deletion/duplication (Smith-Magenis syndrome) region (Figure 5A) [26, 27]. In this locus, all three phenotypes have been associated to varying degrees to one gene, *RAI1* [28]. Curiously, however, only the association signal for EA peaks within this gene, whereas for ASD it unexpectedly maximizes over an adjacent gene. While we believe this tool can be a useful guide to researchers, this example also underscores a limitation of this approach: we cannot provide error rate guarantees at the localized level. Such an analysis will be a target of our future work in this challenging area.

## Data availability statement

The data and code used in this manuscript are available at https://github.com/ryurko/Agglomerative-LD-loci-testing.

## Author contributions statement

R.Y., K.R., B.D., and M.G. conceived the study; R.Y. conducted the analyses; R.Y., K.R., B.D., and M.G. wrote and reviewed the manuscript.

## Supplementary information

### Comparison of GWAS enrichment

We observe that autism spectrum disorder (ASD) GWAS results [2] suffer from weaker power in comparison to schizophrenia (SCZ) [12] and educational attainment (EA) [13] (Supplementary Figure 1). This is likely due to differences in study size: 18,381 cases and 27,969 controls for ASD in comparison to 33,426 cases and 32,541 controls for SCZ and ≈ 1.1 million individuals for EA.

### AdaPT overview

AdaPT is an iterative search procedure for selecting *R* discoveries with guaranteed finite-sample FDR control at target level *α*, under the assumption of independent null *p*-values [8]. We apply AdaPT to a collection of weakly correlated gene/locus-level *p*-values and metadata, (*p_g_, x_g_*)_*g*∈*G*_, testing hypothesis *H_g_* regarding gene/locus’ *g*’s association with the phenotype of interest (e.g. ASD). For each step *t* = 0, 1*, …* in the AdaPT search, we first determine the rejection set 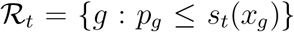, where *s_t_*(*x_g_*) is the rejection threshold at step *t* that is *adaptive* to the metadata *x_g_* (except for the starting threshold *s*_0_ = 0.05). This provides us with both the number of discoveries/rejections 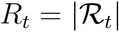, as well as a *pseudo*-estimate for the number of false discoveries *A_t_* = |{*g* : *p_g_* ≥ 1 − *s_t_*(*x_g_*)}|. These quantities are used to estimate the FDP at the current step *t*,

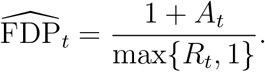

If 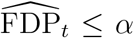, then the AdaPT search ends and the set of discoveries 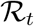 is returned. Otherwise, the rejection threshold is updated by discarding the most likely null element in the current rejection region, as measured by the conditional local false discovery rate (fdr) estimated with an expectation-maximization (EM) algorithm. With the threshold updated, the AdaPT search repeats by estimating FDP and updating the rejection threshold until the target FDR level is reached.

The most critical step in AdaPT involves updating the rejection threshold *s_t_*(*x_i_*) through an EM algorithm to fit a conditional version of the two-groups model [45] to estimate local fdr. We consider the same model form as our previous work [7], where the null p-values are modeled as uniform (*f*_0_(*p*|*x*) ≡ 1) while we model the non-null p-value density with a beta distribution density parametrized by 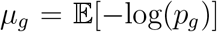. This results in a conditional density for a beta mixture model,

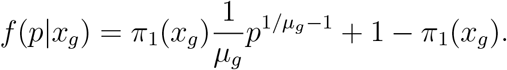

**Supplementary Figure 1:**
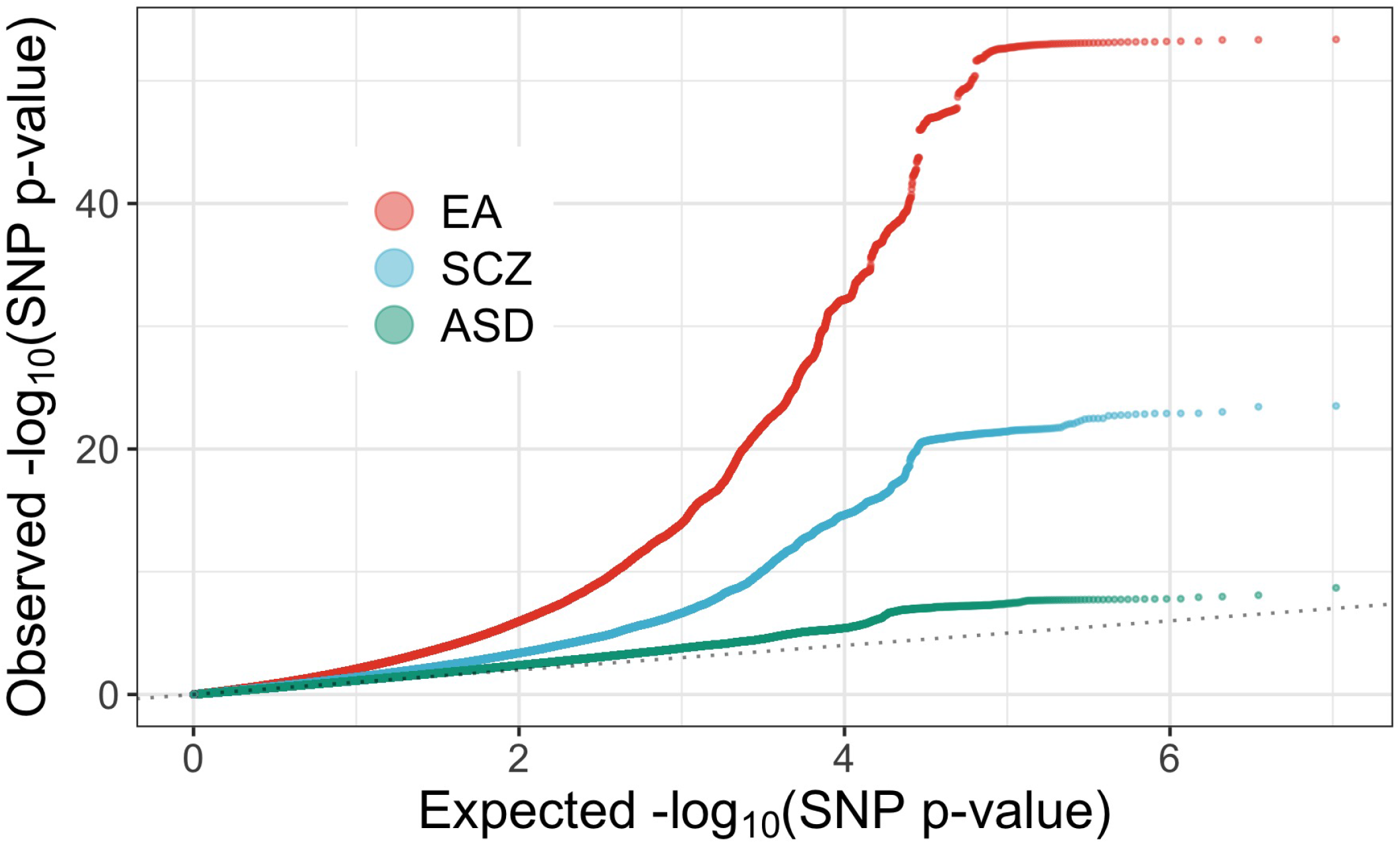
Comparison of SNP-level quantile-quantile plots revealing greater enrichment for SCZ and EA in comparison to ASD. Dotted reference line indicates null, uniform distribution.

In this form, we can model the non-null probability 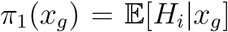 and the non-null effect size 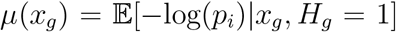 with two separate gradient boosted tree-based models. The XGBoost library [24] provides logistic and Gamma regression implementations which we use for *π*_1_(*x_g_*) and *μ*(*x_g_*) respectively. See our previous work [7] for details about the implementation of the EM algorithm, and [8] for details about updating the AdaPT rejection threshold using *π*_1_(*x_g_*) and *μ*(*x_g_*).

### AdaPT tuning results

In order to avoid overfitting our the non-null probability and effect size models, we first find appropriate settings for the gradient boosted trees (number of trees, learning rate, and maximum depth) using synthetic SCZ *p*-values which mimic ASD *p*-values in the following manner:

1. Sort SCZ and ASD *p*-values: 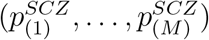 and 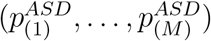
2. Replace SCZ with ASD *p*-values by matching order,

- e.g., replace 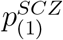 with 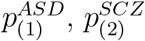 with 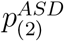, *…*

We then proceed to apply AdaPT using these ASD-aligned SCZ *p*-values, finding appropriate settings in our gradient boosted trees. To avoid overfitting, we consider shallow trees with maximum depth ∈ {1, 2}, while searching over the number of trees *P* ∈ 100*, …, * 450 by increments of fifty and the learning rate *η* ∈ 0.03*, …, * 0.06 by increments of 0.01. We find the combination *P* and *η* yielding the highest number of synthetic SCZ discoveries for both depth values at FDR level *α* = 0.05. The top setting combinations across the considered SNP-to-gene assignment types and *r*^2^ threshold (Supplementary Table 1) were then considered in the AdaPT cross-validation algorithm [7] when applied to the actual ASD, SCZ, and EA *p*-values.

**Supplementary Table 1:** Top boosting settings to consider for number of trees *P* and learning rate *η* by maximum depth and SNP-to-gene assignment type from tuning with synthetic SCZ *p*-values.

| SNP-to-gene assignment | $r^2$ threshold | Depth = 1 | Depth = 2 |
| --- | --- | --- | --- |
| *Positional* | 0.25 | $P = 450, \eta = 0.06$ | $P = 100, \eta = 0.04$ |
| | 0.50 | $P = 150, \eta = 0.06$ | $P = 250, \eta = 0.06$ |
| | 0.75 | $P = 450, \eta = 0.05$ | $P = 200, \eta = 0.05$ |
| *Positional + eSNPs* | 0.25 | $P = 400, \eta = 0.06$ | $P = 250, \eta = 0.05$ |
| | 0.50 | $P = 450, \eta = 0.06$ | $P = 300, \eta = 0.05$ |
| | 0.75 | $P = 450, \eta = 0.05$ | $P = 250, \eta = 0.04$ |

### Measuring AdaPT metadata importance

We examine the variable importance from the gradient boosted trees to provide insight into the relationships between the metadata *x_g_* and measures of phenotypic association, non-null probability *π*_1_(*x_g_*) and non-null effect size *μ*(*x_g_*). However, because AdaPT is an iterative search with several modeling steps, we summarize metadata importance by computing the average importance across the steps in the search. This allows us to compare meta-data importance across phenotypes and positional assignment since the AdaPT searches vary in terms of the number of modeling steps required to reach the target *α* = 0.05 (Supplementary Table 2).

We rank the top five sources of metadata for each phenotype and positional assignment based on the sum of the average importance for the two types of AdaPT models, non-null probability *π*_1_(*x_g_*) and non-null effect size *μ*(*x_g_*). We observe similar rankings in metadata importance between the *Positional + eSNPs* (Figure 3) and *Positional* results (Supplementary Figure 2), with differences between phenotypes such as the increased importance of LOEUF for SCZ and EA in comparison to ASD.

**Supplementary Table 2:** Number of model fitting steps in AdaPT search to reach target FDR level *α* = 0.05, by phenotype and positional assignment.

| Phenotype | *Positional* steps | *Positional + eSNPs* steps |
| --- | --- | --- |
| ASD | 18 | 20 |
| SCZ | 16 | 15 |
| EA | 8 | 8 |

**Supplementary Figure 2:**
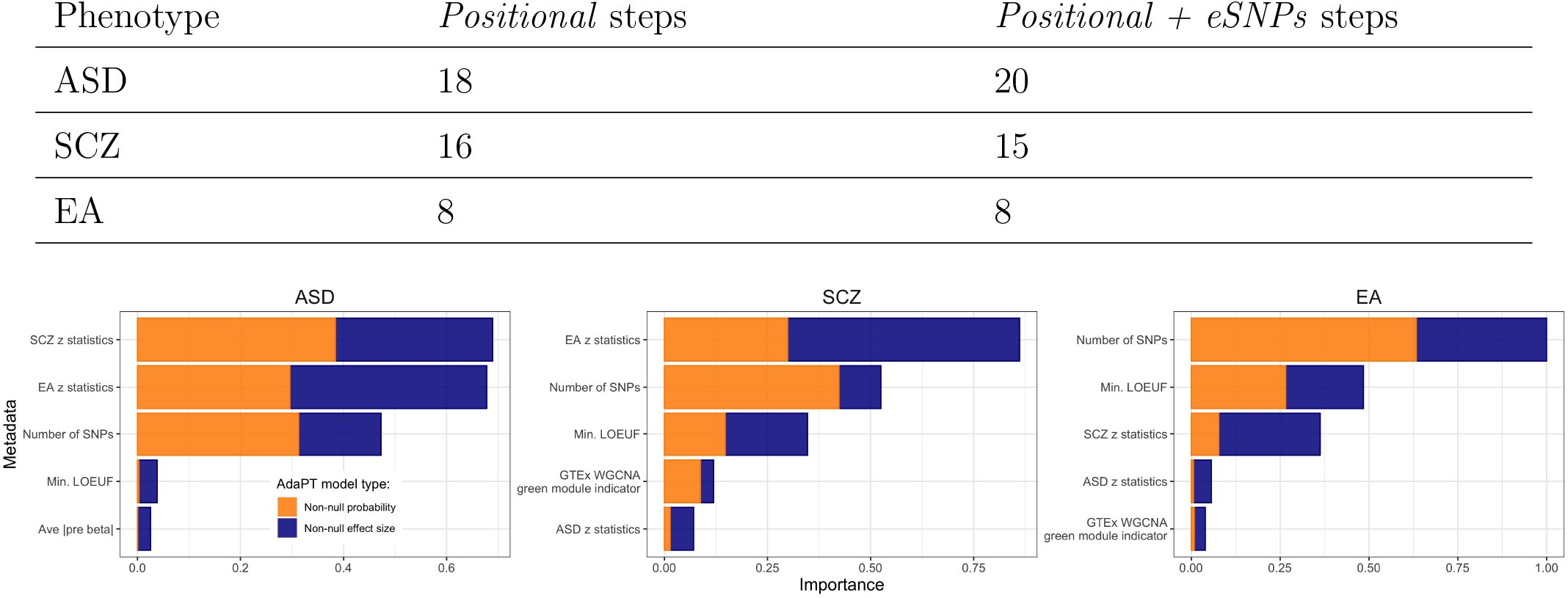
The top five sources of metadata as ranked by XGBoost importance for the *Positional* results by phenotype. Metadata are sorted in order of combined importance, while the color of the bars denote the separate measures of importance for the two AdaPT models: non-null probability (orange) and non-null effect size (blue).

### Supplementary table of results

**Supplementary Table 3:**
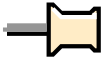
Excel file with six sheets corresponding to the selected genes/loci for each phenotype (ASD, SCZ, or EA) and positional type (Positional + eSNPs or Positional) combination, one sheet with all GSEA results, one sheet with all significant GO terms, and a glossary sheet defining each sheets’ columns.

### Results with LD threshold *r*^2^ ∈ {0.50, 0.75}

We additionally generate the AdaPT: XGBoost and baseline results using genes/loci formed with LD thresholds of *r*^2^ ∈ {0.50, 0.75}, which are not as conservative as the threshold of *r*^2^ = 0.25. Both of these higher threshold values lead to a greater number of genes/loci for testing for both positional types, *Positional* and *Positional + eSNPs* (Figure 2). Ultimately, in terms of the number of genes/loci selected by AdaPT at target FDR level *α* = 0.05, we observe similar results as before (Figure 4A) with increased power from using AdaPT: XGBoost regardless of phenotype with indication that including eSNPs provides an advantage in detecting more signals (Supplementary Figures 3 and 4). As expected, for all three phenotypes, the percentage of genes/loci selected corresponding to unclustered genes increases with the *r*^2^ threshold (Supplementary Table 4).

**Supplementary Table 4:** Comparison of the number (%) of unclustered genes in the AdaPT: XGBoost selected genes/loci as a function of *r*^2^ for each phenotype by positional assignment.

| Positional assignment | $r^2$ | ASD | SCZ | EA |
| --- | --- | --- | --- | --- |
| *Positional* | 0.25 | 77 (17.2%) | 1,439 (30.2%) | 4,991 (42.4%) |
|  | 0.50 | 98 (31.6%) | 2,318 (54.6%) | 6,882 (63.2%) |
|  | 0.75 | 150 (62.2%) | 3,098 (76.5%) | 8,683 (81.9%) |
| *Positional + eSNPs* | 0.25 | 78 (16.1%) | 1,566 (29.3%) | 5,091 (40.5%) |
|  | 0.50 | 112 (28.9%) | 2,306 (50.6%) | 7,054 (61.1%) |
|  | 0.75 | 179 (60.1%) | 3,297 (74.5%) | 9,071 (80.8%) |

### Results per chromosome breakdown

We compare the chromosome breakdown of the selected genes/loci using AdaPT: XGBoost for each phenotype, indicating a greater boost in detecting associations by including eSNPs for ASD in comparison to well-powered SCZ and EA (Supplementary Figure 5). Additionally, for SCZ and EA, the total number of associated genes per chromosome declines strongly and linearly with chromosome size (Supplementary Figures 6 and 7).

**Supplementary Figure 3:**
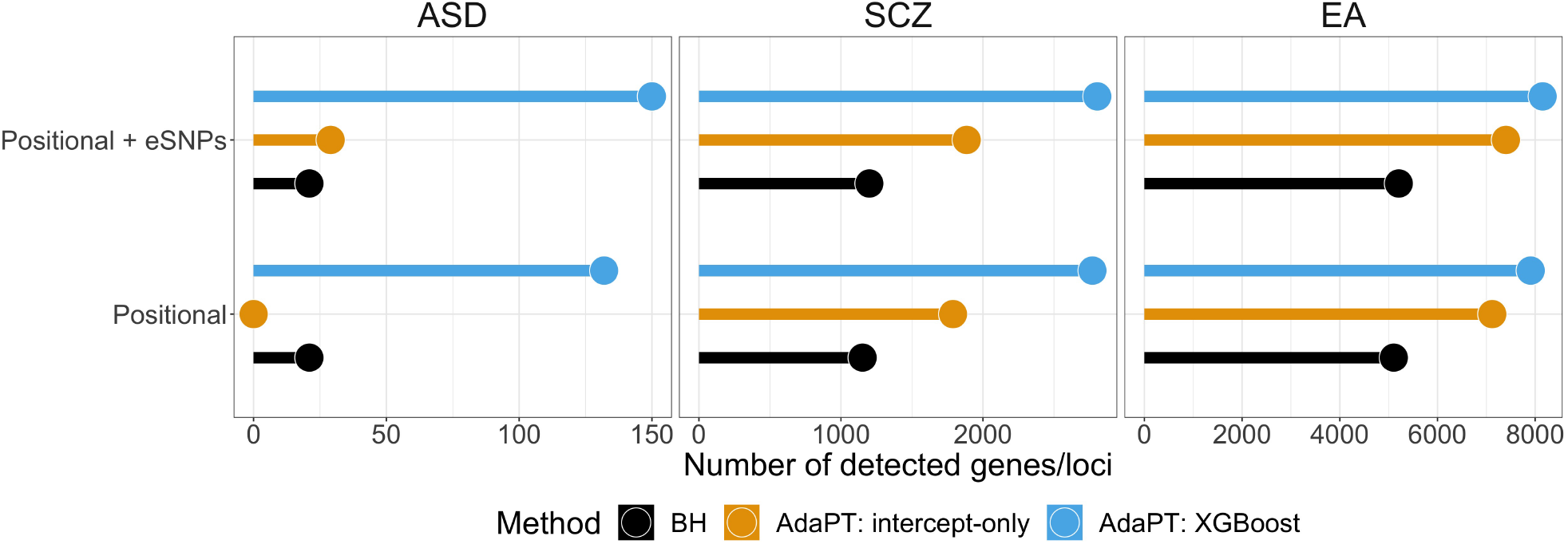
Comparison of the number of discoveries at FDR level *α* = 0.05 for each phenotype (by column), comparing the number of genes/loci returned by the type of SNP-to-gene assignment with LD threshold *r*^2^ = 0.50. AdaPT: XGBoost results are presented in comparison to BH and AdaPT: intercept-only baselines.

**Supplementary Figure 4:**
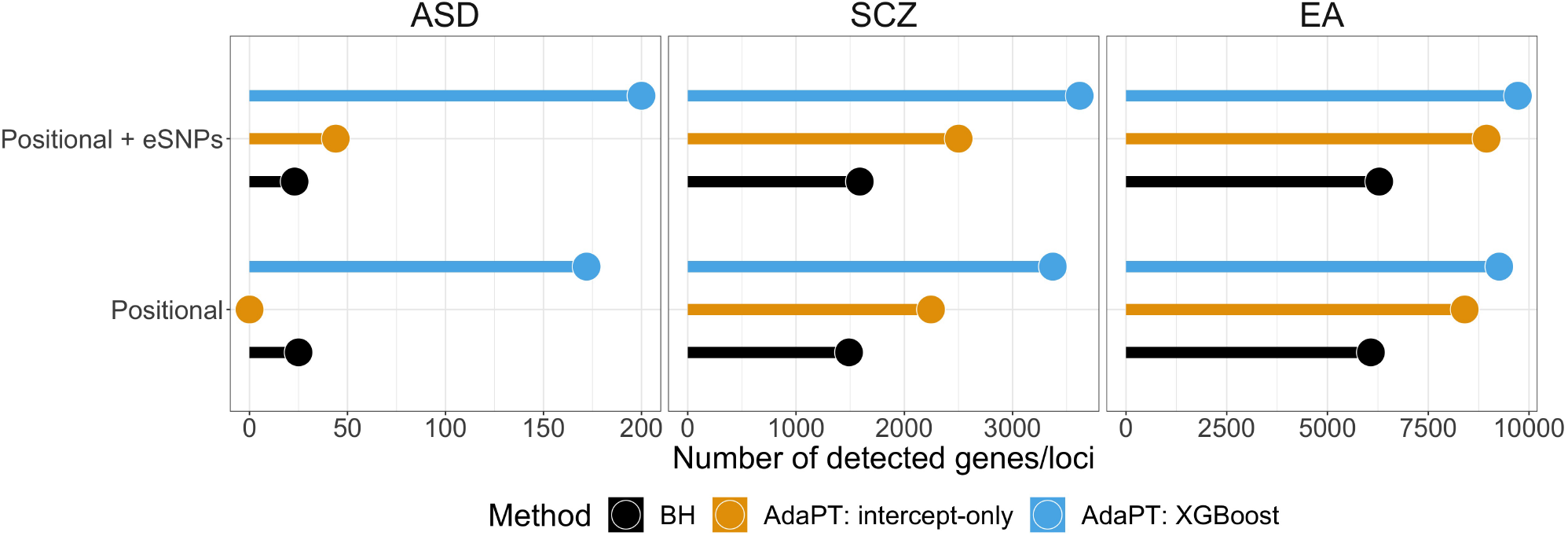
Comparison of the number of discoveries at FDR level *α* = 0.05 for each phenotype (by column), comparing the number of genes/loci returned by the type of SNP-to-gene assignment with LD threshold *r*^2^ = 0.75. AdaPT: XGBoost results are presented in comparison to BH and AdaPT: intercept-only baselines.

**Supplementary Figure 5:**
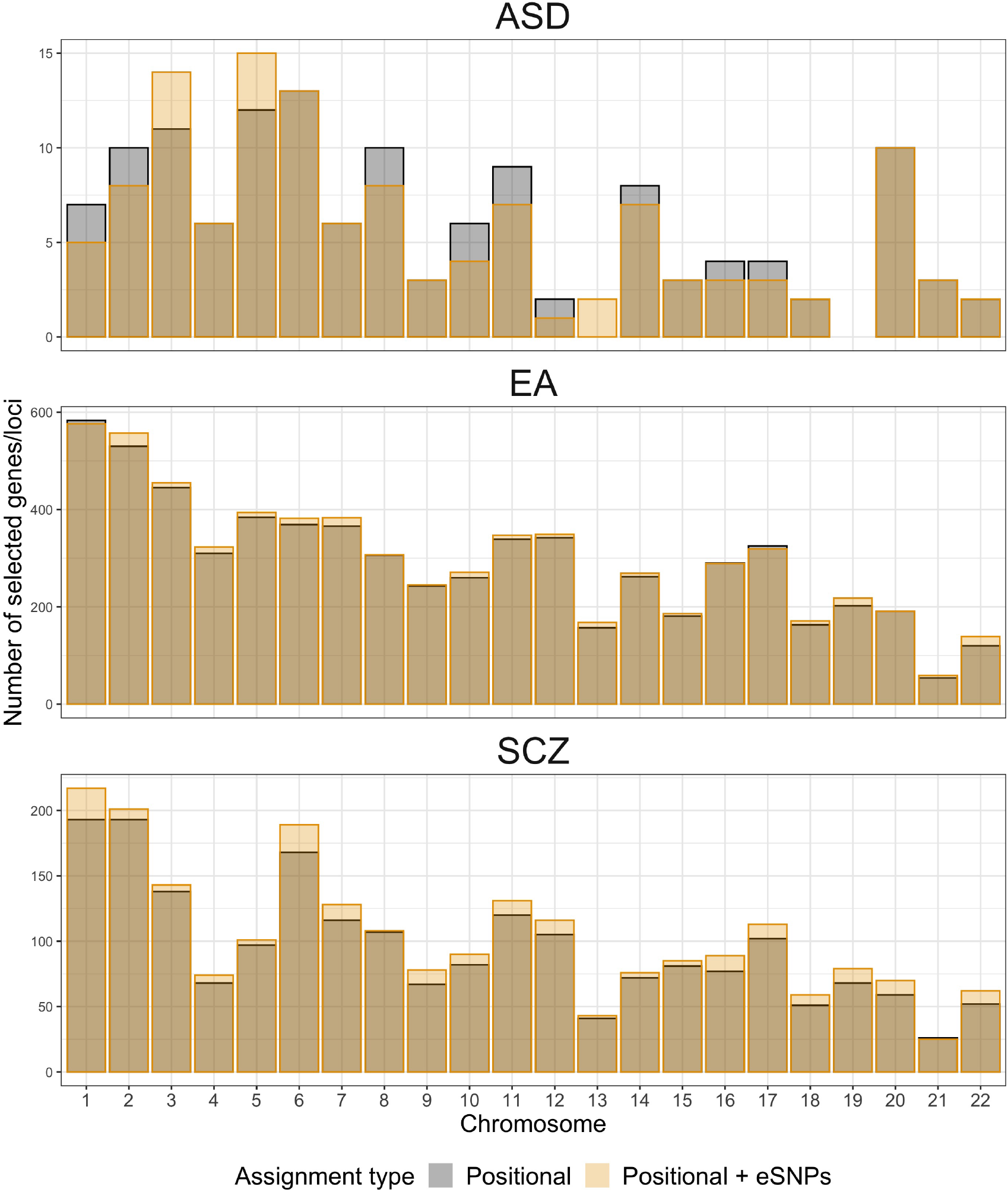
Comparison of the number of selected genes/loci by chromosome in the AdaPT: XGBoost by positional assignment type for each phenotype. The bars are overlaid on top of each other, indicating a higher number of genes/loci are selected at a chromosome using the *Positional + eSNPs* approach when the orange bar height is above the dark gray area, e.g., ASD results for chromosome six.

**Supplementary Figure 6:**
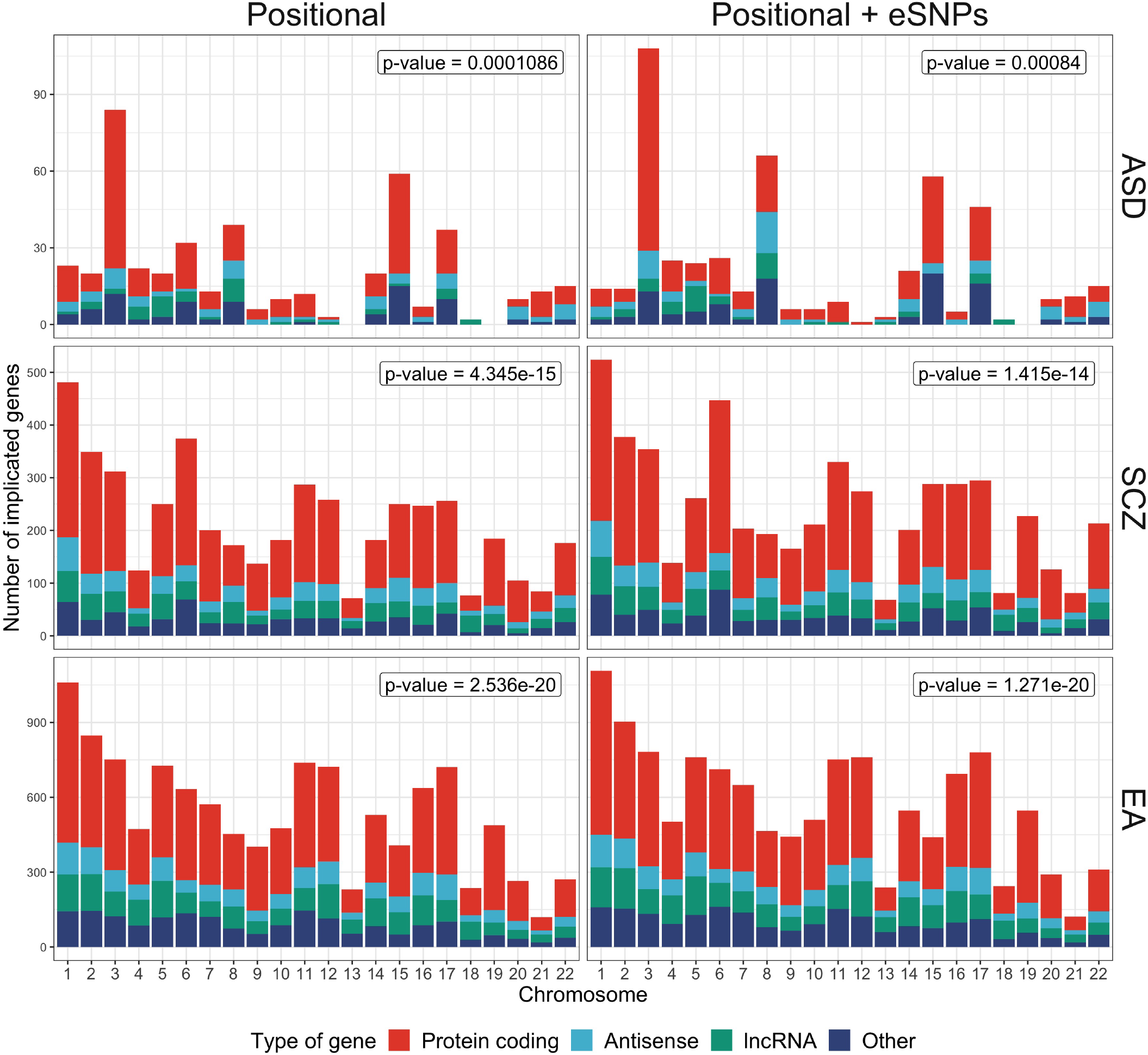
Comparison of the number of implicated genes by chromosome in the AdaPT: XGBoost results for both positional assignment types for each phenotype (colored by type of gene).

**Supplementary Figure 7:**
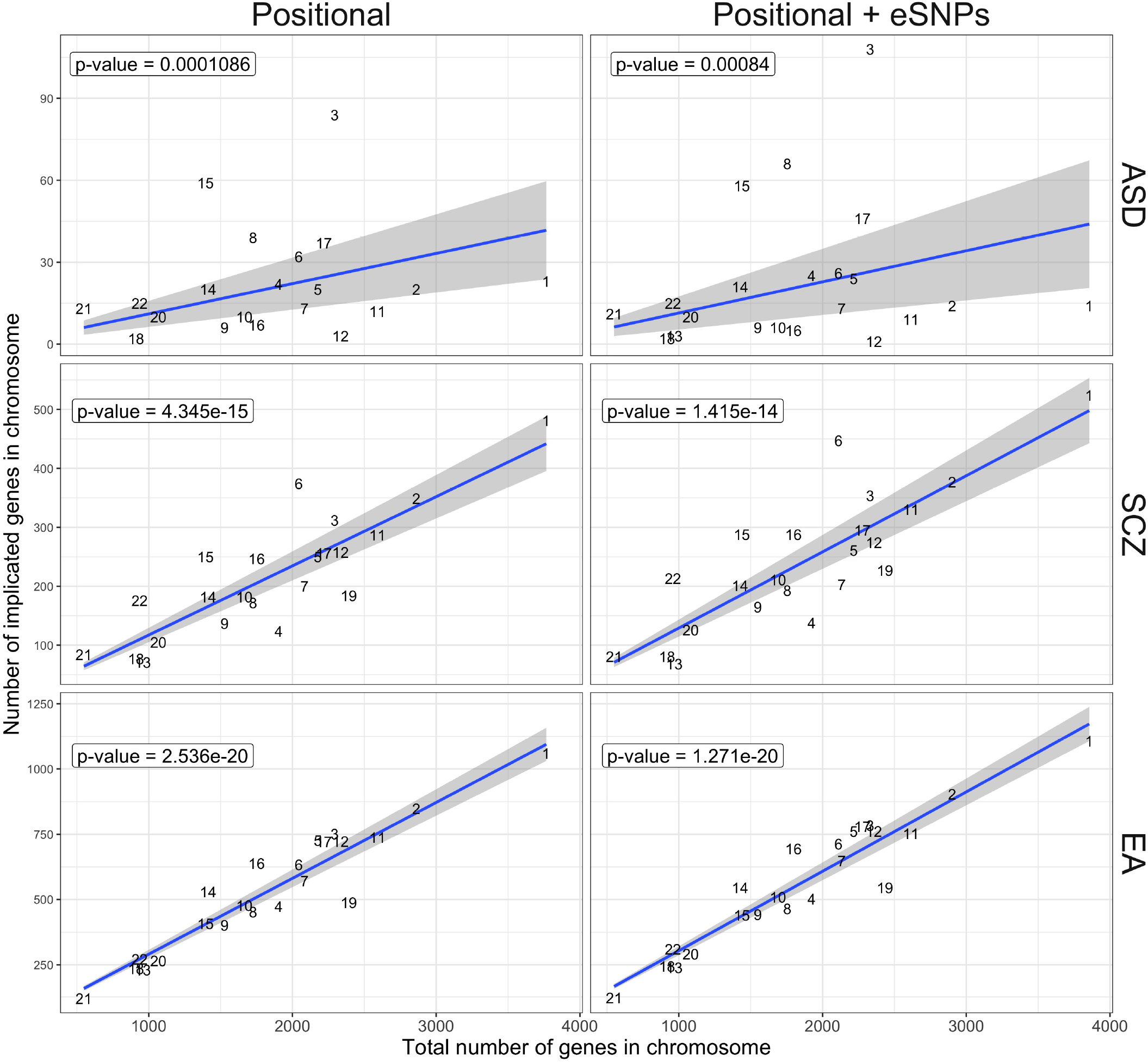
Relationship between the number of implicated genes and total number of genes per chromosome by phenotype and positional assignment. Chromosome points are labeled by their respective number. Blue line indicates regression fit with *p*-values of coefficients labeled in the top-left corners.

### LD locus zoom application

We developed our LD locus zoom application using `R` Shiny [14, 15] and `plotly` [16] to interactively explore localized signal within interesting genes/loci. The genes located within the locus are represented in rectangles denoting their respective start and end positions below the smooth display, arranged by position and size to prevent overlapping. The solid black line denotes the gene/locus-level kernel smoothing for positional SNP signals, including interpolation. For reference, the gray dotted line denotes a point-wise 95^*th*^ percentile of 1,000 simulations for squared null Gaussian random variables, 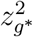 where *z_g*_* ~ Normal(**0**_*g\**_, **Σ**_*g\**_), given the LD structure **Σ**_*g\**_ of the selected LD locus *g\**.

Additionally, we provide several interactive features such as the option to display background kernel smoothing results for SCZ (in red) and EA (in blue), normalized to appear on the same signal scale as ASD, as well as the option to display eSNPs separately as bars (colored to match their associated genes) with bar heights denoting individual eSNP-level signal. Furthermore, one has the option to display gene-level smoothing (Supplementary Figure 8) as well as highlight particular genes and their corresponding eSNP signals with the `plotly` highlighting tool (Supplementary Figure 9). To simplify the visualization of larger loci, we perform the interpolation within subgroup of SNPs that are formed if they are separated by more than five percent of the loci size (using single-linkage clustering), e.g. three subgroups of SNPs with interpolation performed separately for LD locus in chromosome 17 1 Mb inversion region (Supplementary Figure 9).

Within the application, there are three tabs: (1) *ASD results*, (2) *Upload results*, and (3) *Description*. The *ASD results* tab includes the LD locus visualization for our *Positional* and *Positional + eSNPs* results, with the ability to select different genes/loci to display as well as export the image as an SVG file. Furthermore, the tables of the selected LD locus’ corresponding genes and SNPs can be downloaded as CSV or Excel files (via the *Genes* and *SNPs* tabs respectively). The gene and SNP tables also include urls to their respective pages in the GWAS catalog [46].

Additionally, users can use the *Upload results* tab to import datasets to generate the same type of kernel smoothing visualization. Additional information about the interactive features of the application are available in the *Description* tab. The LD locus zoom application can be accessed here https://ron-yurko.shinyapps.io/ld_locus_zoom/.

### Enrichment analysis

A primary motivation for gene-based analyses is to garner insight into the biological mechanisms underlying the phenotype by evaluating the set of genes associated with it. We implement two approaches: (1) FUMA GENE2FUNC tool [38] for gene ontology (GO) enrichment analysis of the AdaPT: XGBoost genes/loci, and (2) gene-set enrichment analysis (GSEA) [39] to test if different sets of genes are enriched in a ranked gene list. In both approaches, we address the confounding effect of gene/locus size as larger genes are more likely to be associated (Supplementary Figure 10).

First, we use the FUMA GENE2FUNC tool for GO enrichment analysis of the implicated genes in the AdaPT: XGBoost genes/loci. Rather than include all implicated genes in the associated loci for the FUMA enrichment analysis, we used the kernel smoothing results to identify *signal* genes. Specifically, we only use genes with kernel smoothing signals above the point-wise 95th percentile of null simulations and any gene with at least one intergenic eSNP displaying a large marginal effect size (i.e., z statistic ≥ 1.96). Then to address the confounder of gene size, we find non-implicated genes to match the *signal* genes with respect to gene size using the `optmatch` [47] package in `R` [14]. Due to the difference in number of *signal* genes for the phenotypes, we find the twenty, two, and the single closest matching non-implicated genes for each *Positional + eSNPs signal* gene with ASD, SCZ, and EA respectively. Using the list of matched non-implicated background genes, we then do not observe any GO enrichment terms for the ASD genes. In comparison, the SCZ (EA) genes display enrichment for 698 (114) terms in biological processes, 163 (64) terms in cellular components, and 103 (27) in molecular function. Using REVIGO [44] to summarize these terms, they highlight neuron project, synaptic function, cell adhesion, cell cycle, chromosome organization, and and many more (Supplementary Figures 11 and 12).

Next, we performed GSEA for five different sets of genes/loci to test for enrichment at the top of the list of genes/loci ranked by their one-sided z statistics:

1. 18,008 brain expressed genes from SynGO [48],
2. 1,232 synaptic genes from SynGO [48],
3. 815 epigenetic genes from EpiFactors [49],
4. 1,819 transcription factors compiled from the SeqQC project [50], and
5. 102 ASD risk genes identified based on *de novo* and case-control variation [40].

We collapsed these five lists of genes into their respective genes/loci based on LD induced correlation for each positional type, indicating that a locus is a member of a list if at least one of its genes is a member of the list (Supplementary Table 5).

**Supplementary Table 5:**
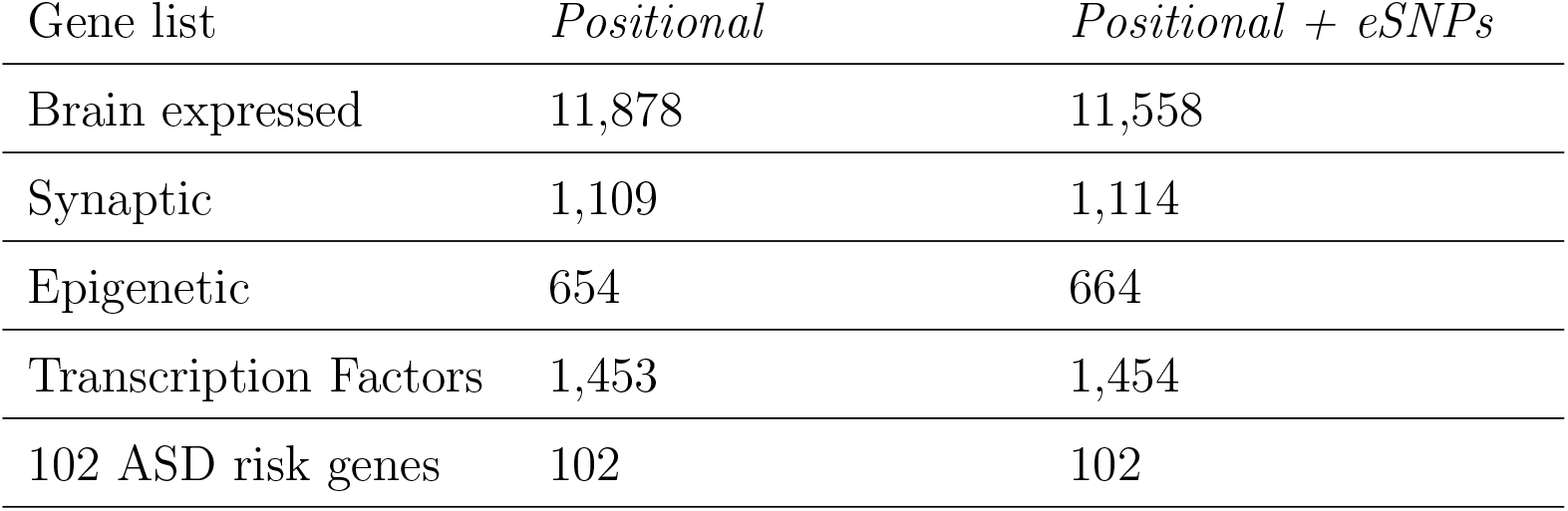
Comparison of the number of genes/loci in each of the five considered list of genes for GSEA by positional assignment.

| Gene list | *Positional* | *Positional + eSNPs* |
| --- | --- | --- |
| Brain expressed | 11,878 | 11,558 |
| Synaptic | 1,109 | 1,114 |
| Epigenetic | 654 | 664 |
| Transcription Factors | 1,453 | 1,454 |
| 102 ASD risk genes | 102 | 102 |

To address the confounder of gene/locus size, we compute versions of the z statistics that are adjusted for the gene/locus’ size. Specifically, we regress out the effect of the log(size) on the z statistics for each phenotype (Supplementary Figure 10) and use the adjusted z statistics for GSEA. We use the fgsea implementation in R [51], with 10,000 permutations to compute GSEA *p*-values, to test if any of the five lists are enriched at the top of the genes/loci ordered by the size-adjusted z statistics for all three phenotypes. We observe that both synaptic and epigenetic genes/loci are enriched for both positional types using the size-adjusted ASD z statistics based on Benjamini-Hochberg (BH) [52] adjusted *p*-values at FDR level *α* = 0.05 (Figure 6 and Supplementary Figure 13). In comparison, we observe that all five gene sets, for both positional types, are enriched for both SCZ and EA size-adjusted z statistics (all of their respective BH adjusted *p*-values were equal to 1 / 10,001, i.e., their observed enrichment scores were more extreme than the 10,000 simulations).

**Supplementary Figure 8:**
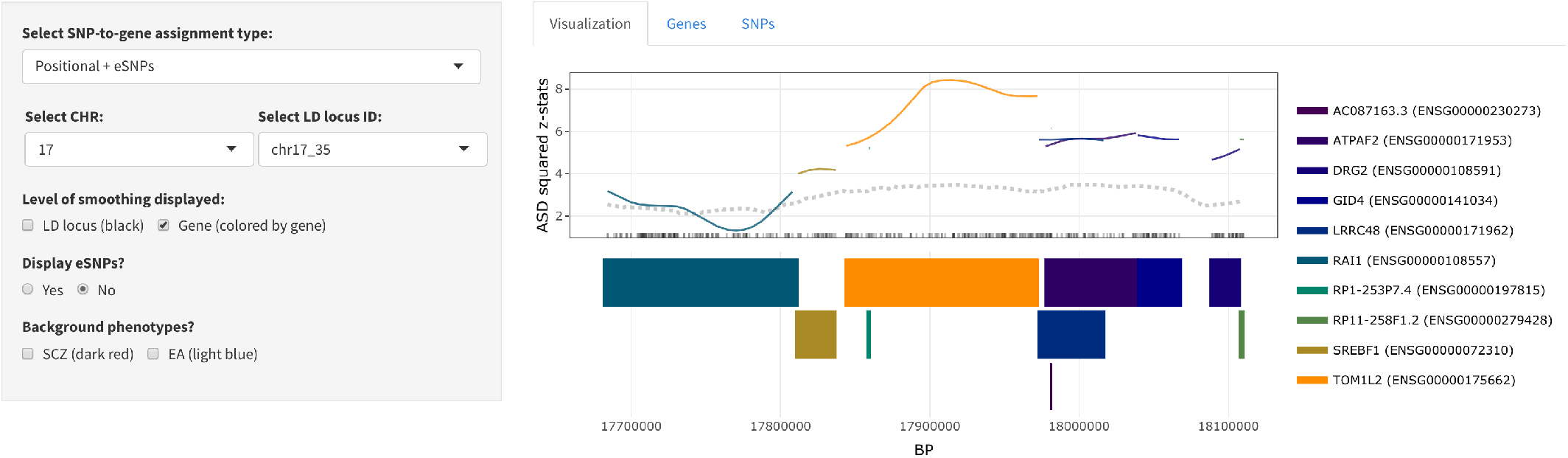
Gene-level ASD kernel smoothing for the chromosome 17 LD locus in the 3.7 Mb deletion region. Line colors match the associated genes displayed below the x-axis. The gray dotted line indicates point-wise 95^*th*^ percentile for null simulations.

**Supplementary Figure 9:**
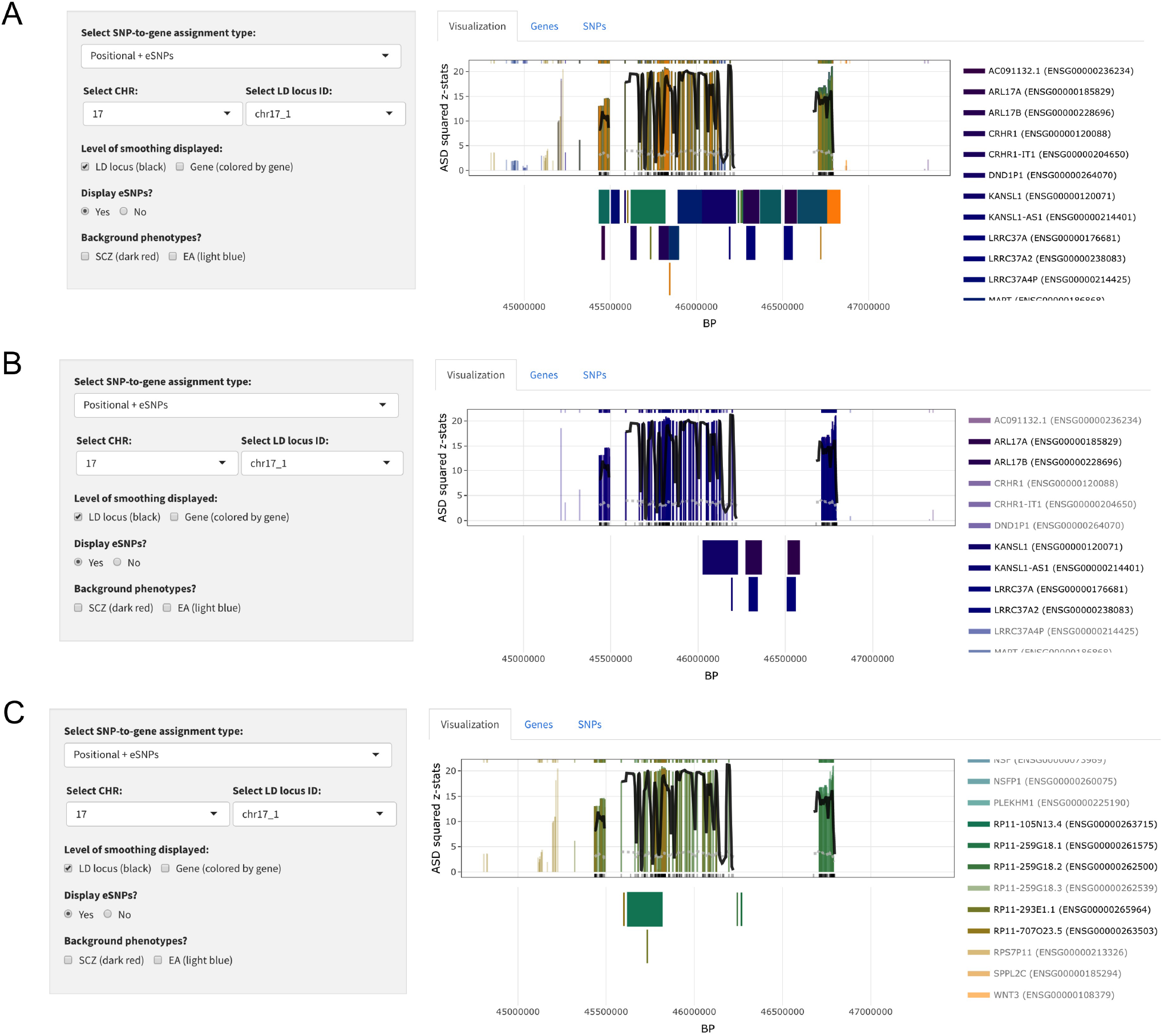
Zoom display for LD locus in chromosome 17 1 Mb inversion region. *(A)* Display with all genes and their associated eSNPs. *(B)* and *(C)* Display two separate subsets of genes and their associated eSNPs.

**Supplementary Figure 10:**
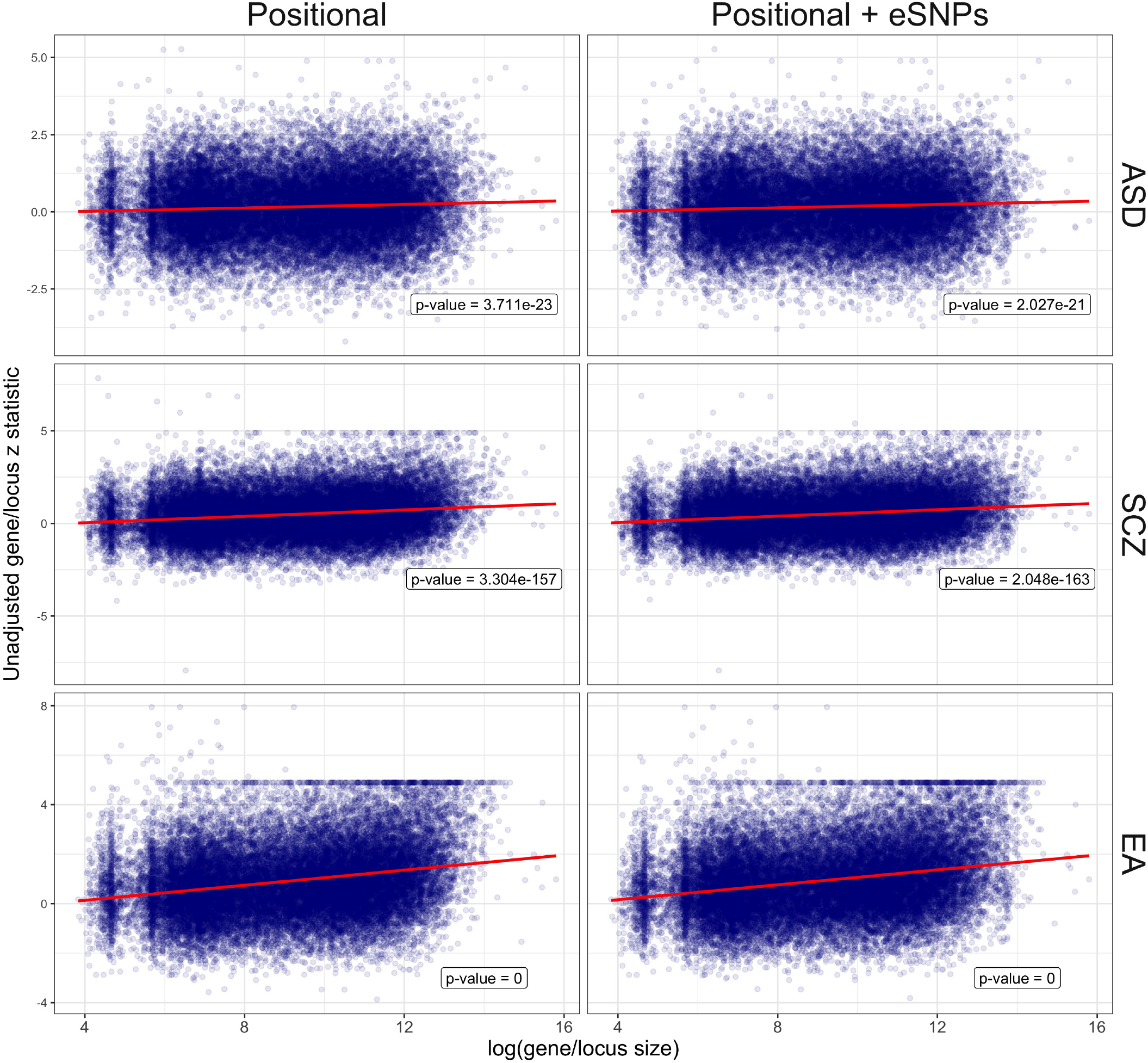
Relationship between unadjusted z statistics and log(gene/locus size) by phenotype and positional assignment. Red line indicates regression fit with *p*-values of coefficients labeled in the bottom-right corners.

**Supplementary Figure 11:**
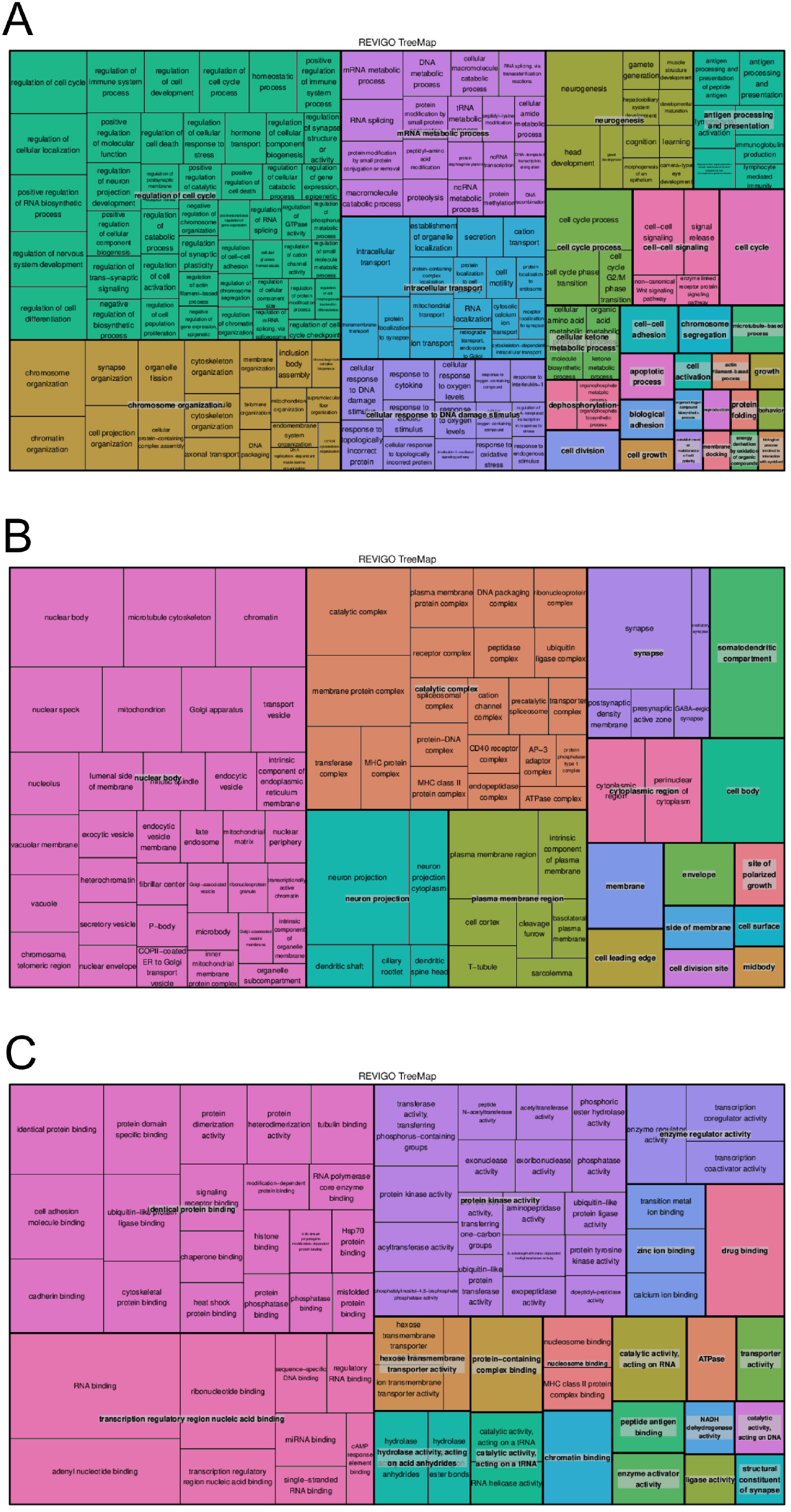
Treemap display of GO *(A)* biological processes, *(B)* cellular components, and *(C)* molecular function for SCZ *Positional + eSNPs* results using *signal* genes with size-matched list of background genes.

**Supplementary Figure 12:**
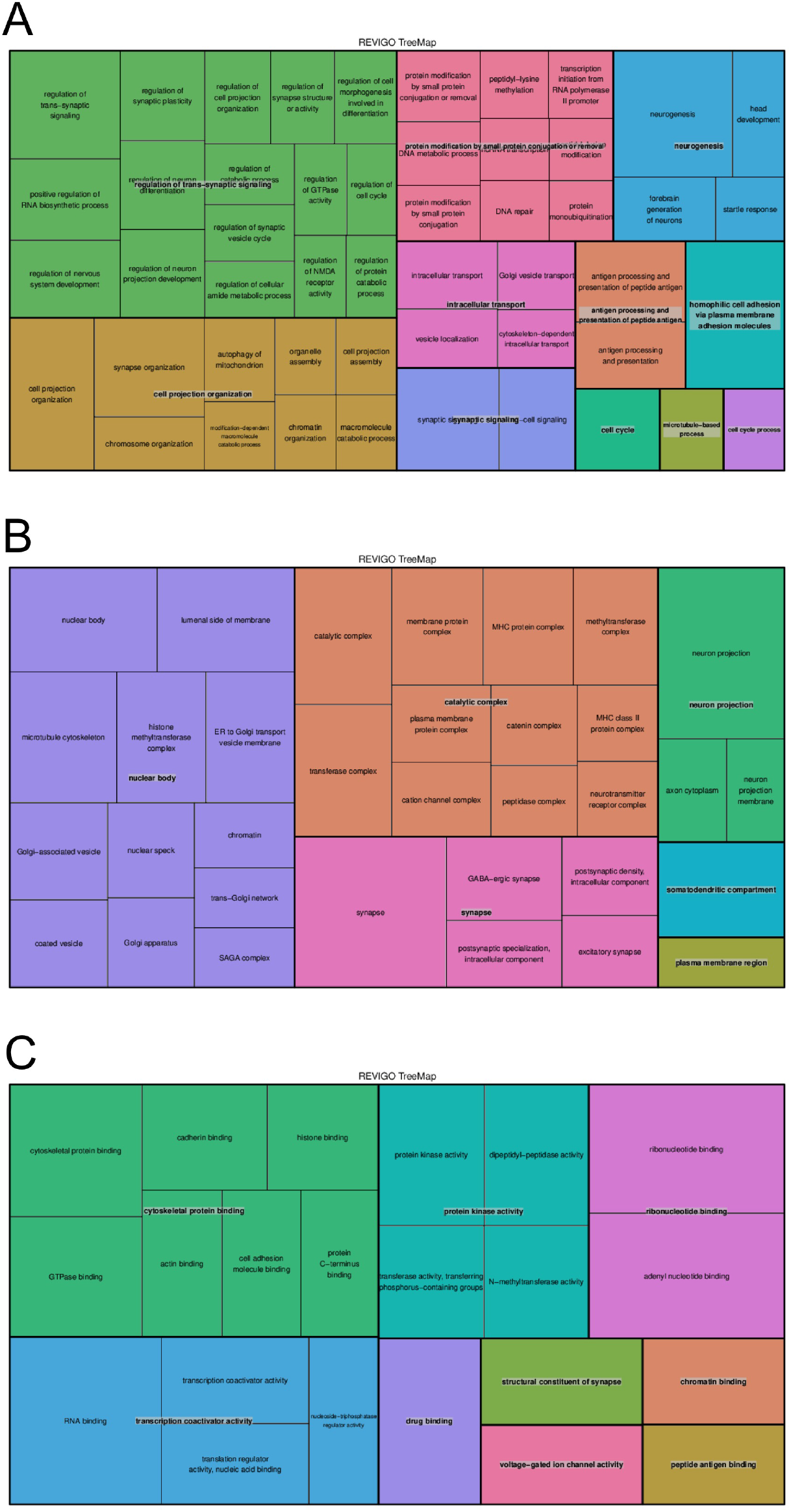
Treemap display of GO *(A)* biological processes, *(B)* cellular components, and *(C)* molecular function for EA *Positional + eSNPs* results using *signal* genes with size-matched list of background genes.

**Supplementary Figure 13:**
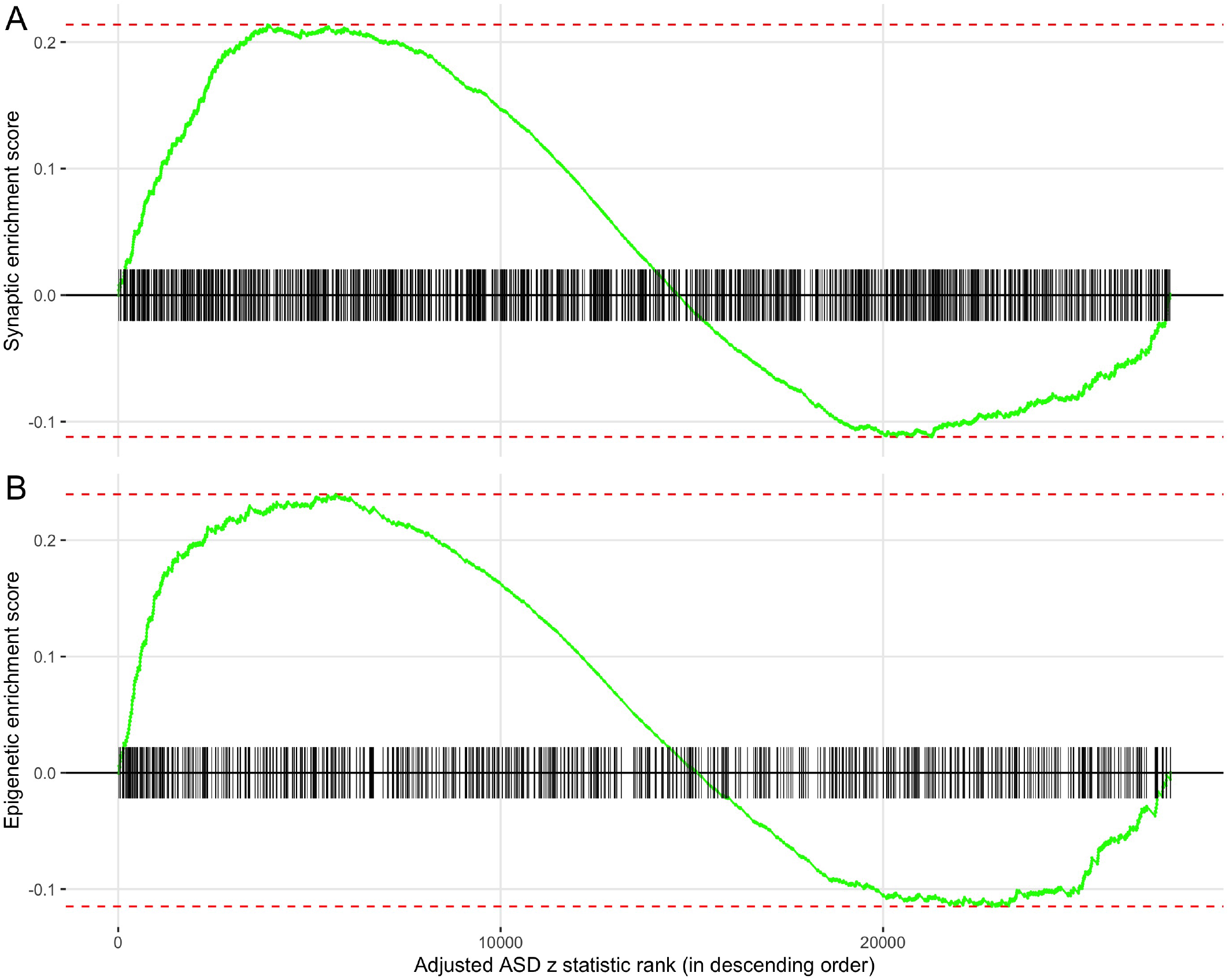
Enrichment score for *(A)* synaptic and *(B)* epigenetic genes/loci with tick marks denoting gene/locus rank based on *Positional* ASD z statistics, adjusted for size, in descending order.

